# An indigenous *Saccharomyces uvarum* population with high genetic diversity dominates uninoculated Chardonnay fermentations at a Canadian winery

**DOI:** 10.1101/838268

**Authors:** Garrett C. McCarthy, Sydney C. Morgan, Jonathan T. Martiniuk, Brianne L. Newman, Vivien Measday, Daniel M. Durall

**Affiliations:** Irving K. Barber School of Arts and Sciences, Department of Biology, The University of British Columbia, Kelowna, British Columbia, Canada; Wine Research Centre, Faculty of Land and Food Systems, The University of British Columbia, Vancouver, British Columbia, Canada

**Keywords:** *Saccharomyces uvarum*, spontaneous fermentation, microsatellite, indigenous yeast, phylogenetic tree, low-temperature fermentation

## Abstract

*Saccharomyces cerevisiae* is the primary yeast species responsible for most fermentations in winemaking. However, other yeasts, including *Saccharomyces uvarum*, have occasionally been found conducting commercial fermentations around the world. *S. uvarum* is typically associated with white wine fermentations in cool-climate wine regions, and has been identified as the dominant yeast in fermentations from France, Hungary, northern Italy, and, recently, Canada. However, little is known about how the origin and genetic diversity of the Canadian *S. uvarum* population relates to strains from other parts of the world. In this study, a highly diverse *S. uvarum* population was found dominating uninoculated commercial fermentations of Chardonnay grapes sourced from two different vineyards. Most of the strains identified were found to be genetically distinct from *S. uvarum* strains isolated around the world. Of the 106 strains of *S. uvarum* identified in this study, four played a dominant role in the fermentations, with some strains predominating in the fermentations from one vineyard over the other. Furthermore, two of the dominant strains were previously identified as dominant strains in uninoculated Chardonnay fermentations at the same winery two years earlier, providing evidence for a winery-resident population of indigenous *S. uvarum*. This research provides a valuable insight into the diversity and persistence of non-commercial *S. uvarum* strains in North America, and provides a stepping stone for future work into the enological potential of an alternative *Saccharomyces* yeast species.

## Introduction

Modern winemaking often involves the inoculation of grape must with one or more commercial yeast strains, usually belonging to the dominant winemaking yeast species *Saccharomyces cerevisiae*. However, single-strain fermentations have been shown to produce less complex wines than fermentations conducted by multiple yeast species and strains [1–3]. Furthermore, the commercial *S. cerevisiae* strains used in these inoculated single-strain fermentations may be aggressively competitive towards indigenous yeasts. The loss of indigenous yeast strains during the winemaking process can result in the reduction of regional character, because the local consortium of microorganisms can contribute to the expression of *terroir* in wine [3,4]. In recent years, many winemakers have become interested in conducting uninoculated (spontaneous) fermentations in an attempt to encourage a diversity of yeast species and strains to participate in alcoholic fermentation. Although vineyard-derived yeasts were originally thought to be the ones conducting uninoculated fermentations, a growing body of evidence has shown that uninoculated fermentations are actually conducted by winery-resident yeast strains [5–8]. These yeasts may be of commercial or indigenous origin, but have established themselves as residents of the winery environment, and are capable of entering and conducting fermentations over multiple vintages.

Although most uninoculated fermentations at commercial wineries are conducted by strains of *S. cerevisiae* [5–8], some wineries contain established populations of *Saccharomyces uvarum* that are able to conduct and complete alcoholic fermentation [9–12]. *S. uvarum* belongs to the *Saccharomyces sensu stricto* clade, and is the furthest relative from *S. cerevisiae* within this clade [13]. The Holarctic *S. uvarum* population, which originated in the northern hemisphere [14–16], includes both natural *S. uvarum* strains isolated from *Quercus* (oak) trees as well as all industrial strains isolated from cider, beer, and wine fermentations. It is believed that this population evolved alongside other *Saccharomyces* species, as 95% of Holarctic *S. uvarum* strains possess introgressed regions from *Saccharomyces eubayanus, Saccharomyces kudriavzevii*, and *S. cerevisiae* throughout their genome [14,15]. However, the history of *S. uvarum* research is difficult to trace, because *S. uvarum* has had many names, and has even shared names with now distinct species. In the past, *S. uvarum* has been referred to as *Saccharomyces bayanus* var. *uvarum* [10,11,17,18] or simply *Saccharomyces bayanus* [19,20]. To complicate matters, many commercial *S. cerevisiae* strains have been marketed incorrectly as strains of *S. bayanus* [21]. However, *S. uvarum* is now known to be a pure species, distinct from *S. bayanus*, which itself is a hybrid of the pure species *S. uvarum* and *S. eubayanus* [13,15,22,23].

*S. uvarum* is a cryotolerant yeast usually found in association with white wine fermentations in cool-climate wine regions [9,10,12,24], but has also been associated with cider production [19,20] and some traditional fermentations [25,26]. During fermentation, *S. uvarum* produces lower levels of ethanol, acetic acid, and acetaldehyde, and higher levels of glycerol, succinic acid, malic acid, isoamyl alcohol, isobutanol, and ethyl acetate, as compared to *S. cerevisiae* [27–30]. Additionally, because of its ability to conduct fermentation at lower temperatures, *S. uvarum* may produce wines with more balanced aroma profiles [31]. However, few studies have been conducted to investigate the origins, genetic diversity, and enological potential of this yeast, thus more research is needed on this topic.

The overall objective of this study was to identify the presence and genetic diversity of *S. uvarum* strains conducting uninoculated fermentations at a commercial winery in the Okanagan Valley wine region of British Columbia, Canada, and place this population within the context of *S. uvarum* strains from around the world. Two years before this current study was conducted, a highly diverse, indigenous population of *S. uvarum* was identified at this same commercial winery [12]. We were interested in investigating the persistence of *S. uvarum* strains in the winery environment over multiple vintages. A secondary objective involved sampling fermentations containing grape must from two different vineyards, to investigate whether the geographical origin or chemistry of the grapes played a role in determining the fungal communities and *S. uvarum* populations present in the fermentations.

## Materials and methods

### Experimental design and sampling

This study was conducted in association with Mission Hill Family Estate Winery during the 2017 vintage. The fungal diversity and community composition of grapes from two different vineyards were followed from the vineyard into the winery and throughout alcoholic fermentation. Samples for high-throughput amplicon sequencing (Illumina MiSeq) were taken from grapes in the vineyard just prior to harvest, as well as at four stages of fermentation in the winery. Samples for *Saccharomyces uvarum* population analysis were taken at three stages of alcoholic fermentation.

#### Vineyard grape samples

Two Chardonnay vineyards managed by Mission Hill Family Estate Winery in the Okanagan Valley of British Columbia, Canada, were selected for this study (Vineyard 2 and Vineyard 8). The exact locations of these vineyards, along with the dates of grape sample collection, can be viewed in Table 1. Both vineyards had been herbicide-free since 2016 (2017 was the second herbicide-free vintage), and were transitioning from conventional to organic viticulture practices.

**Table 1.** Summary of vineyards used in this study.

| Vineyard name | Closest municipality | Latitude | Longitude | Sample collection | Date of commercial harvest |
| --- | --- | --- | --- | --- | --- |
| Vineyard 2 | Osoyoos, BC | 49.000603 | -119.418712 | 2017-09-04 | 2017-09-05 |
| Vineyard 8 | Oliver, BC | 49.221125 | -119.559834 | 2017-09-18 | 2017-09-19 |

Each vineyard was divided into six sampling sections (achieved by dividing the number of rows in the vineyard by six), and was further sub-divided into potential sampling sites within each section. Each post in the vineyard was planted approximately 15 feet (4.57 m) apart, with five vines planted in between each post (within each panel). A potential sampling site was defined as three consecutive panels (equalling approximately 15 vines), and one sampling site was randomly selected within each sampling section, for a total of six samples per vineyard (S1 Fig). The following restrictions were placed on randomization: sampling occurred at least three posts inward from the edge of each vineyard, and at least two rows away from neighbouring vineyard blocks, to minimize the potential for contamination from nearby roads or other grape varietals. Sampling was performed by aseptically collecting one cluster from each vine in the sampling site, on both sides of the row, for a total of 30 clusters per sample. The 30 clusters from each sampling site were placed into a sterile bag (one bag per sampling site, six bags per vineyard) and transported on ice back to the laboratory at The University of British Columbia (Kelowna, BC, Canada) for same-day processing. Each bag containing grape clusters was then gently crushed and homogenized by hand for 10 min, after which time 2 mL samples of the juice were collected and frozen at −80°C to await further processing for high-throughput amplicon sequencing.

#### Winery samples

The winery portion of this study was conducted at Mission Hill Family Estate Winery, a large commercial winery in British Columbia, Canada that conducts both inoculated and uninoculated (spontaneous) fermentations of many different grape varietals and makes wines in many different styles. The Chardonnay grapes for this study were sourced from two vineyards (Vineyard 2 and Vineyard 8), as described above. The must from each vineyard was crushed and pressed into large stainless steel tanks to undergo a cold settling period before being transferred into 285 L stainless steel barrels (La Garde, SML Stainless Steel, Québec, Canada). Must from Vineyard 2 spent two days in the cold settling tank, while must from Vineyard 8 spent eight days in the cold settling tank. Six stainless steel barrels were used in this experiment: three contained must exclusively from Vineyard 2, and three contained must exclusively from Vineyard 8. Because Vineyard 2 is located further south than Vineyard 8 (Table 1), the grapes from this vineyard ripened earlier, and were harvested before those from Vineyard 8, in order to achieve similar sugar concentrations at crush. Although the fermentations from each vineyard began at different times, they did overlap in the same cellar for 20 days during sampling. The cellar where the fermentations were conducted was maintained at 12 °C.

Samples were taken at four stages of uninoculated (spontaneous) alcoholic fermentation as defined by their sugar concentration (approximated by Brix level): cold settling (22 °Brix, prior to the start of AF), early (14-18 °Brix), mid (6-10 °Brix), and late (2 ± 0.1 °Brix). To each barrel, 20 mg/L total sulfur dioxide was added in the form of potassium metabisulfite (K_2_S_2_O_5_) during the cold settling stage, 250 ppm Lallemand^®^ Fermaid K complex yeast nutrient was added between the cold settling and early stages, and 100 ppm Laffort^®^ THIAZOTE mineral nutrient was added between the early and mid stages. Samples were collected in sterile 50 mL centrifuge tubes and were transported on ice to the laboratory at The University of British Columbia (Kelowna, BC, Canada) for processing. Subsamples (2 mL) were washed and frozen at −80 °C for high-throughput amplicon sequencing, while other subsamples were processed immediately for culture-dependent *S. uvarum* strain typing.

### Chemical analyses

Chemical parameters for the grape must and wine were taken at the cold settling and late stages, respectively. Yeast assimilable nitrogen (YAN), titratable acidity, volatile acidity, malic acid, pH, residual sugar, ethanol content, fructose, and glucose were measured using a WineScan^TM^ wine analyzer (Foss, Hillerød, Denmark).

### Saccharomyces uvarum strain typing

#### Colony isolation and DNA extraction

Samples from the early, mid, and late stages of fermentation were serially-diluted and plated onto YEPD media (1% (w/w) yeast extract (BD Bacto^TM^, Sparks, MD, USA), 1% (w/w) bacterial peptone (HiMedia Labs, Mumbai, India), 2% (w/w) dextrose (Fisher Chemical, Fair Lawn, NJ, USA), 2% (w/w) agar (VWR, Solon, OH, USA)), with the addition of 0.01 % (v/v) chloramphenicol (Sigma-Aldrich, St. Louis, MO, USA) to prevent bacterial growth, and 0.015 % (v/v) biphenyl (Sigma-Aldrich, St. Louis, MO, USA) to prevent filamentous fungal growth [9,32–34]. Plates were incubated at 28 °C for 48 h and stored at 4 °C. Plates containing 30-300 colonies were selected for colony isolation and DNA extraction. Grape samples from the vineyard and the cold settling stage samples were not analyzed for *S. uvarum* strains, because *Saccharomyces* yeasts are rarely found in must before the onset of alcoholic fermentation [35], and are present on healthy grapes in very low abundance [36].

From the early, mid, and late stage plates, 47 single yeast colonies were isolated onto Wallerstein Nutrient media (WLN) (Sigma-Aldrich, St. Louis, MO, USA), a differential medium used to identify non-*S. cerevisiae* yeasts. Two controls were used to distinguish between *S. cerevisiae* and *S. uvarum* colonies: Lalvin^®^ BA11 (Lallemand, Montreal, QC, Canada) and CBS 7001 (Westerdijk Fungal Biodiversity Institute, Utrecht, Netherlands), respectively. The WLN plates were incubated at 28 °C for 48 h, and then stored at 4 °C. *S. cerevisiae* isolates appeared as medium-sized, cream-coloured colonies, while *S. uvarum* isolates appeared as small, forest green colonies. Only the *S. uvarum* isolates were selected for strain identification. DNA was extracted from each yeast isolate by performing a water DNA extraction, as described previously [5].

#### Multiplex PCR and fragment analysis

*S. uvarum* strain typing was performed as previously described [12] using a multiplex PCR reaction targeting 11 microsatellite loci that had been selected from two previous studies: L1, L2, L3, L4, L8, L9 [37], NB1, NB4, NB8, and NB9 [24]. Briefly, multiplex PCR was performed on the extracted DNA from each isolate and submitted to Fragment Analysis and DNA Sequencing Services at the University of British Columbia (Kelowna, BC, Canada) for fragment analysis on a 3130xl DNA sequencer (Applied Biosystems, Foster City, CA, USA). GeneMapper 4.0 software (Applied Biosystems, Foster City, CA, USA) was used to determine the fragment size at each locus, and the resulting multilocus genotype of each isolate was compared to that of the others using Bruvo’s genetic distance [38]. Bruvo’s distance was calculated in RStudio (version 3.5.2) using the ‘poppr’ package (version 2.8.1) [39,40], and applying a genetic distance threshold of 0.3 for isolate comparison. Yeast isolates obtained in this study were compared to a database containing 150 *S. uvarum* strains identified during the 2015 vintage at the same winery [12], as well as 12 international *S. uvarum* strains from around the world (Table 2).

**Table 2.**
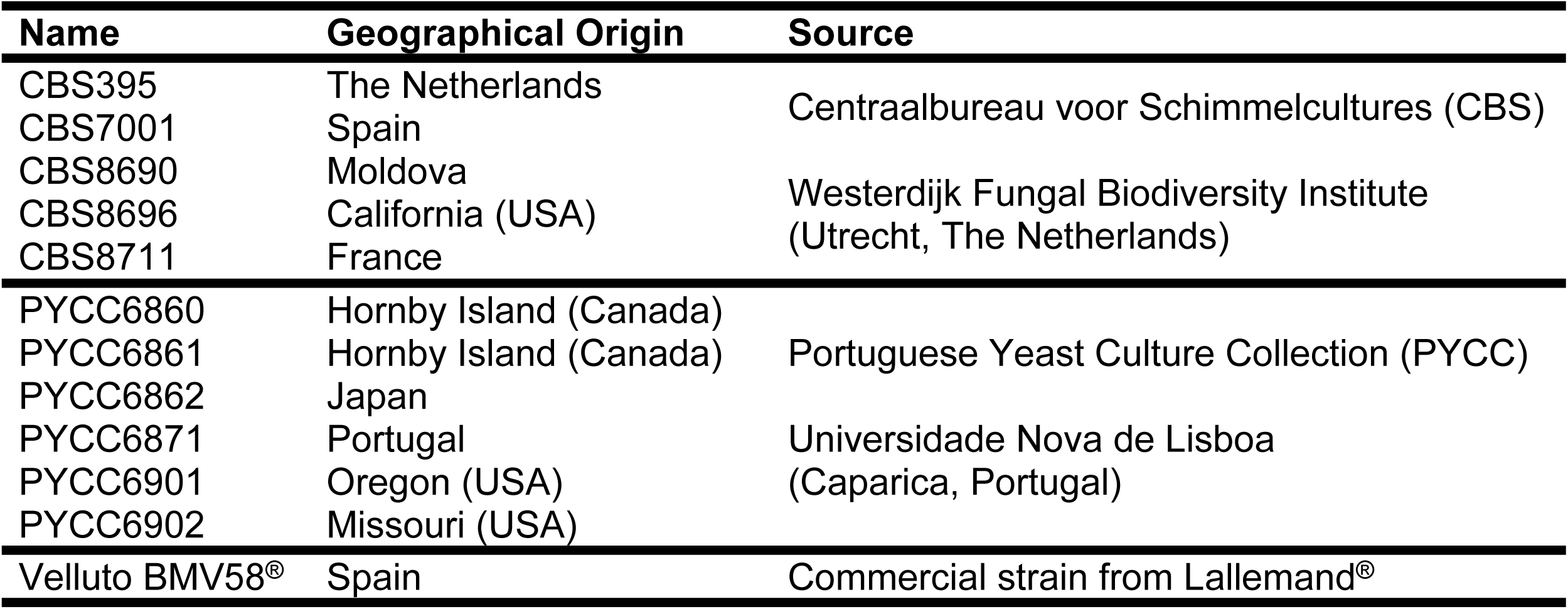
Names, geographical origins, and sources of international *Saccharomyces uvarum* strains used for comparison in this study.

Isolates that only partially amplified were re-run and were subsequently excluded from analysis upon a second failure. Isolates that did not amplify were considered to be non-*S. uvarum* yeasts and were also excluded from analysis. Although 47 yeast isolates were originally selected from each sample, not all belonged to *S. uvarum*, and after strain identification, each sample was rarefied to 32 *S. uvarum* isolates.

#### Yeast selection for phylogenetic tree construction

An unrooted, neighbour-joining phylogenetic tree was generated to compare the genetic relatedness of the significant *S. uvarum* strains identified in Chardonnay fermentations in the Okanagan Valley winemaking region of Canada (in both the 2015 and 2017 vintages), relative to *S. uvarum* strains identified in other regions of the world. Twelve international *S. uvarum* strains were selected for this purpose (Table 2). Significant *S. uvarum* strains are defined as those that represented > 1% of all the yeast isolates strain-typed in each vintage. The phylogenetic tree was generated in RStudio (version 3.5.2) using the ‘ape’ package (version 5.2) [41] as well as the “find.clusters” function in the ‘adegenet’ package (version 2.0.1) [42], with *k*-means successive clustering for statistical grouping of subpopulations in the tree.

### High-throughput amplicon sequencing

#### Sample treatment and DNA extraction

Samples for high-throughput amplicon sequencing (Illumina MiSeq) were taken from grapes in the vineyard (grapes), as well as from four stages of fermentation in the winery (cold settling, early, mid, and late).

Samples (previously frozen at −80 °C) were thawed on ice, and then washed before total DNA was extracted following a modified protocol from a previously-published study [43]. Samples were pelleted by centrifugation at 13,200 rpm for 5 min. The supernatant was discarded and the pellet was re-suspended in 1 mL chilled 1× phosphate-buffered saline (Sigma-Aldrich, St. Louis, MO, USA), then centrifuged again at 13,200 rpm for 5 min. The supernatant was discarded and the pellet was re-suspended in 500 µL of 50 mM ethylenediaminetetraacetic acid (EDTA, pH 8.0) (Invitrogen, Grand Island, NY, USA). Samples were mixed by pipetting up and down five times, and the entire sample was transferred to a FastPrep tube (MP Biomedicals, Santa Ana, CA, USA) containing 200 mg of 0.5 mm glass disruptor beads (Scientific Industries, Bohemia, NY, USA). The FastPrep tubes were placed into a Vortex-Genie 2 Digital bead beater (Scientific Industries, Bohemia, NY, USA) for two rounds of 2.5 min (30/s), separated by 1 min on ice. Aliquots of 500 µL Nuclei Lysis solution (Fisher, Hampton, VA, USA) were added to the FastPrep tubes and lysed in the bead beater for 1 min (30/s). Samples were then incubated for 10 min at 95 °C and then centrifuged at 13,200 rpm for 5 min. To an autoclaved 2 mL microcentrifuge tube (VWR, Radnor, PA, USA), 500 µL of the supernatant was added, followed by 250 µL Protein Precipitation solution (Fisher, Hampton, VA, USA). Samples were then vortexed lightly and kept at room temperature (22 °C) for 15 min. Samples were then centrifuged at 13,200 rpm for 5 min, and 500 µL of the supernatant was transferred to new 2 mL microcentrifuge tubes containing 75 µL 20% (v/v) polyvinylpyrrolidone (PVP) solution (Sigma-Aldrich, St. Louis, MO, USA). Samples were pulse-vortexed for 10-20 sec and centrifuged at 13,200 rpm for 10 min. Supernatant (500 µL) was transferred to a new 2 mL microcentrifuge tube containing 300 µL chilled 2-propanol (Sigma-Aldrich, St. Louis, MO, USA). Tubes were inverted to mix several times and left to sit at room temperature for 15 min. Samples were then centrifuged at 13,200 rpm for 2 min and the supernatant was discarded. The pellet was re-suspended in 1 mL chilled 100% ethanol (Commercial Alcohols, Brampton, ON, Canada), centrifuged at 13,200 for 2 min, and the entire supernatant was carefully discarded. Samples were left open in a biosafety cabinet for a maximum of 30 minutes to ensure complete evaporation of the ethanol. The samples were re-suspended in 50 µL of 10 mM TE buffer (Invitrogen, Grand Island, NY, USA), and frozen at −80 °C. A positive control, containing pure *S. cerevisiae* cells, and a negative control, containing only molecular-grade water, were also subjected to the same DNA extraction protocol, and were included during library preparation and Illumina sequencing.

#### Illumina MiSeq library preparation

Sample library preparation used a two-step PCR procedure consisting of ‘amplicon’ and ‘index’ PCR reactions, as described previously [44]. Briefly, amplicon PCR was performed by amplifying the ITS1 region of the rRNA gene using BITS and B58S3 primers [45] with CS1 and CS2 linker sequences, respectively. Index PCR primers contained Illumina MiSeq adapter sequences, unique eight nucleotide barcodes, 9-12 bp heterogeneity spacers, and CS1/CS2 linker sequences. After both PCR reactions, samples were submitted to the IBEST Genomics Resources Core at the University of Idaho (Moscow, ID, USA) for quantification, normalization, pooling, and sequencing. Paired-end sequencing (300 bp) was performed on an Illumina MiSeq Desktop Sequencer (Illumina Inc., San Diego, CA, USA).

#### Illumina MiSeq data processing

Illumina MiSeq data processing was performed using both R (version 3.5.1) and the open-source bioinformatics pipeline Quantitative Insights Into Microbial Ecology (QIIME2 version 2018.11) [46]. In R, sequences were denoised using the ‘dada2’ package (version 1.8) [47], as well as the ‘ShortRead’ (version 1.36.1) [48] and ‘Biostrings’ (version 2.46.0) packages. All forward and reverse primer sequences had been removed from the 5’ end of the sequences by IBEST at the University of Idaho before being returned, but because some ITS sequences are likely to be shorter than 300 bp, it is possible that these sequences contain nucleotides from the opposite primer, which required removal before further processing using Cutadapt [49]. After all primer sequences had been removed, sequences were filtered and trimmed using the “filterandTrim” function in the ‘dada2’ package. Any sequence containing an N was removed, as well as any sequence shorter than 50 bp. The maximum number of expected errors allowed in any sequence was set to 2. Because the ITS1 region in fungi is highly variable [50,51], trimming all sequences to the same length can reduce the diversity of the identified community and can even remove sequences with true lengths shorter than the specified truncation length. For this reason, sequences were not trimmed to a consistent length. Forward and reverse reads were merged using the “mergePairs” function, a sequence table was constructed with the “makeSequenceTable,” and chimeras were removed with the “removeBimeraDenovo” function. The representative sequence table was converted to a Fasta file before being transferred from R to the QIIME2 pipeline to complete analysis.

In QIIME2, sequences underwent paired-end alignment using MAFFT [52], and a phylogenetic tree with a mid-point root was created using FastTree 2 [53]. Sequence variants were classified to the species level (if possible) using a dynamic (97-99%) threshold classifier made with the UNITE (version 8.0) database [54]. Sequence variants that could not be classified to the order level or lower and those that appeared with a total frequency of < 100 sequences were excluded from analysis. Samples were rarefied to 20,000 sequences before being exported from QIIME2 for statistical analysis and visualization. Two samples did not meet the applied threshold of 20,000 sequences, and were therefore removed from analysis: one sample was a cold settling sample from the Vineyard 8 fermentations, and the other was a grape sample from Vineyard 2. In the UNITE (version 8.0) database, *S. uvarum* is incorrectly classified as *S. bayanus*, because in the past both species were considered synonymous. Based on our culture-dependent data, we are confident that sequences classified in the UNITE database as *S. bayanus* belong to *S. uvarum*, and we have accordingly re-named all our sequences identified as *S. bayanus* to *S. uvarum*.

Software packages, versions, parameters, and primer sequences used in this study can be viewed at https://osf.io/j7rx8/. The QIIME2 artifact can be viewed at https://view.qiime2.org. For raw sequence data, please visit https://qiita.ucsd.edu and search for Study 12837 (EBI submission ID to be updated).

### Statistical analysis

The chemical parameters of the grape must and wine from each vineyard were statistically compared by performing a 1-way analysis of variance (ANOVA), using the “aov” function in RStudio (version 3.5.2) (α = 0.05). Each parameter was evaluated separately. Normality was not assessed due to the low number of samples. The assumption of homogeneity of variance was tested using the “leveneTest” function in the ‘Rcmdr’ package (version 2.5-1). Levene’s test indicated no violation in the assumption of homogeneity of variance in any chemical parameter (data not shown). Because only two groups were being compared, a post-hoc test was not necessary.

Simpson’s Index of Diversity (1 − *D*) was calculated using the “diversity” function in the ‘vegan’ package (version 2.5-1) in RStudio (version 3.5.2) and reported ± the standard error of the mean (SEM). Diversity was evaluated at four stages of fermentation for the fungal communities, and at three stages of fermentation for the *S. uvarum* strains.

Because the fermentations from both vineyards did not occur simultaneously (Vineyard 8 was harvested 14 days after Vineyard 2), and because the Vineyard 8 must spent longer in the large settling tank, it is possible that the differences in fungal communities and *S. uvarum* populations observed in this study are not solely a result of differences in vineyard geography. Therefore, we have chosen not to perform inferential statistics on these data, but have instead chosen to visually display the results in order to enable a discussion of the differences and trends observed. The relative abundance of fungi and *S. uvarum* strains was visualized by creating stacked bar charts using GraphPad Prism (version 8.2.1) software (La Jolla, CA, USA). A principal coordinates analysis (PCoA) was generated using “vegdist” and “wcmdscale” functions in the ‘vegan’ (version 2.5-3) package in RStudio (version 3.5.2), in order to visualise the spatial distribution of the *S. uvarum* population among samples and treatments. All raw data and R scripts used in the preparation of this manuscript can be viewed at https://osf.io/j7rx8/.

## Results and discussion

### Fermentation progression and wine chemistry

The total time from harvest of the Chardonnay grapes to the end of alcoholic fermentation was similar for the must from both vineyards: the must from Vineyard 2 completed fermentation 40 days after harvest, and the must from Vineyard 8 completed fermentation 37 days after harvest. However, the time the must spent in the settling tank before being transferred to barrels for fermentation differed between the two vineyard treatments. Must from Vineyard 2 spent only two days in the large stainless steel settling tank, while must from Vineyard 8 spent eight days, due to a higher proportion of solid material that required a longer settling time.

Sugar concentration at crush was similar between the two vineyards (Table 3). Although a significant difference was found between the vineyards for sugar concentration (°Brix), this is likely because of the low variation observed within each treatment. It is unlikely that a difference of less than 1 °Brix would have any biologically relevant effects on the microbial composition of the wine. pH was higher in the must from Vineyard 2, while yeast assimilable nitrogen, titratable acidity, and malic acid were all significantly higher in the must from Vineyard 8.

**Table 3.** Chemical parameters of stainless steel barrel-fermented Chardonnay wines sourced from two vineyards. Samples were taken at the cold settling stage (prior to the start of alcoholic fermentation) and at the late state (towards the end of alcoholic fermentation). Values are the mean ± SEM (n = 3 per treatment). An asterisk next to the chemical parameter indicates a significant difference between the two vineyards. Each chemical parameter was evaluated separately within each stage.

| Stage | Chemical Parameter | Vineyard 2 | Vineyard 8 |
| --- | --- | --- | --- |
| Cold settling | pH* | 3.37 ± 0.006 | 3.31 ± 0.003 |
| | Residual sugar ($^{\circ}$Brix)* | 22.3 ± 0.0 | 22.0 ± 0.03 |
| | Yeast assimilable nitrogen (mg/L)* | 77.7 ± 5.6 | 110.7 ± 3.9 |
| | Titratable acidity (g/L)* | 5.07 ± 0.03 | 7.20 ± 0.0 |
| | Malic acid (g/L)* | 2.20 ± 0.0 | 3.47 ± 0.03 |
| Late | pH | 3.37 ± 0.01 | 3.41 ± 0.01 |
| | Titratable acidity (g/L)* | 7.20 ± 0.2 | 7.83 ± 0.03 |
| | Malic acid (g/L)* | 1.97 ± 0.03 | 2.40 ± 0.06 |
| | Volatile acidity (g/L)* | 0.47 ± 0.003 | 0.38 ± 0.001 |
| | Ethanol (% v/v) | 11.7 ± 0.1 | 11.3 ± 0.1 |
| | Glucose (g/L) | 1.07 ± 0.1 | 1.37 ± 0.3 |
| | Fructose(g/L) | 28.9 ± 1 | 27.1 ± 2 |

By the late stage of alcoholic fermentation, there was no significant difference between the wines from the two vineyards in terms of pH, ethanol content, or glucose and fructose concentration (Table 3). Very little glucose remained in the wine by the end of alcoholic fermentation (< 2 g/L), while close to 30 g/L fructose remained unfermented. *S. uvarum*, which dominated the fermentations from both vineyards, is a glucophilic yeast [29], and the residual fructose observed in the late stage of fermentation suggests that this yeast is less adept at fermenting fructose than *S. cerevisiae*. This result is supported by previous research [55].

By the late stage, titratable acidity and malic acid were still significantly higher in the wines from Vineyard 8 than the wines from Vineyard 2 (Table 3). However, the change in malic acid from the cold settling to the late stage was also much greater in the wines from Vineyard 8. In the wines from Vineyard 2, the malic acid concentration decreased from 2.20 g/L to 1.97 g/L, a decrease of approximately 10%. Meanwhile, in the wines from Vineyard 8, the malic acid concentration changed from 3.47 g/L to 2.40 g/L, a decrease of approximately 30%. This could suggest a more significant presence of malic acid-degrading bacteria in the must from Vineyard 8. Indeed, a previous study conducted at this same winery with grapes from Vineyard 8 identified *Tatumella* spp. in barrels that did not receive SO_2_ [12]. These bacteria were thought to originate in the vineyard, as they were present in vineyard samples, but seemed to be sensitive to SO_2_; the bacteria were not able to persist in treatments that received 40 mg/L SO_2_ at crush. When they did persist, however, the malic acid was almost completely degraded by the end of alcoholic fermentation. It is possible that the addition in this current study of only 20 mg/L SO_2_ allowed a portion of these bacteria to survive in the must and perform a partial degradation of malic acid. Unfortunately, we did not identify the bacterial community in this current study, so more research is needed to confirm this hypothesis.

Volatile acidity was found to be significantly higher in the wines from Vineyard 2 (Table 3). However, neither vineyard produced wines with unacceptable levels of volatile acidity, and all the wines in this study contained volatile acidity levels that were below the detection threshold of 0.7 g/L [56].

### Fungal communities

#### Fungal community diversity

Fungal species diversity was highest in the grape and cold settling samples for both vineyards, and decreased for the early, mid, and late stages of fermentation (Table 4). A decrease in the diversity of fungal taxa is expected at the onset of alcoholic fermentation, because most yeasts and fungi present on grapes are non-fermentative, and are either killed by the presence of ethanol or absence of oxygen, or out-competed for space and nutrients by the more dominant yeasts [35,57]. Diversity was constant throughout the three alcoholic fermentation stages (early, mid, and late) for both vineyard treatments, but the wines from Vineyard 8 had a consistently higher diversity than the wines from Vineyard 2.

**Table 4.** Fungal species diversity, measured as Simpson’s Index of Diversity (1 − *D*), of stainless steel barrel-fermented Chardonnay sourced from two different vineyards. Diversity ± SEM was measured from grape samples taken in the vineyard (grapes), as well as at four stages of fermentation in the winery (cold settling, early, mid, and late). Vineyard 2 grape sample values are the means ± SEM of five replicates, and Vineyard 8 grape sample values are the means ± SEM of 6 replicates. All winery fermentation stages have three reported replicates, with the exception of the cold settling stage from the Vineyard 8 fermentations, which contained two.

| Sample | Vineyard 2 | Vineyard 8 |
| --- | --- | --- |
| Grapes | 0.85 ± 0.03 | 0.69 ± 0.05 |
| Cold settling | 0.64 ± 0.03 | 0.87 ± 0.02 |
| Early | 0.064 ± 0.005 | 0.21 ± 0.09 |
| Mid | 0.061 ± 0.009 | 0.31 ± 0.04 |
| Late | 0.061 ± 0.009 | 0.30 ± 0.1 |

#### Fungal community composition

In total, 194 fungal taxa were identified in this study, of which 11 achieved ≥ 10% relative abundance in at least two samples (S1 Table). The fungal community composition from each of the two vineyards differed markedly at every sampling stage (Fig 1). Interestingly, the fungal communities of the grape and cold settling samples within each vineyard treatment were also very different. The cold settling samples were taken after the grapes had been harvested, crushed, and processed, so at least some of the differences observed between these stages can be attributed to contact with winery equipment. In the Vineyard 2 treatment, the most abundant fungi in the grape samples were *Alternaria* sp., *Mycosphaerella tassiana*, and *Epicoccum nigrum*, while the most abundant fungi in the cold settling samples were *Aspergillus niger*, *Aureobasidium pullulans*, and *Penicillium* sp. In the Vineyard 8 treatment, the most abundant fungi in the grape samples were *Erysiphe necator* and *Mycosphaerella tassiana*, while the most abundant fungi in the cold settling samples were *Candida* sp., *Penicillium* sp., *S. uvarum*, and *Hanseniaspora osmophila*.

**Fig 1.**
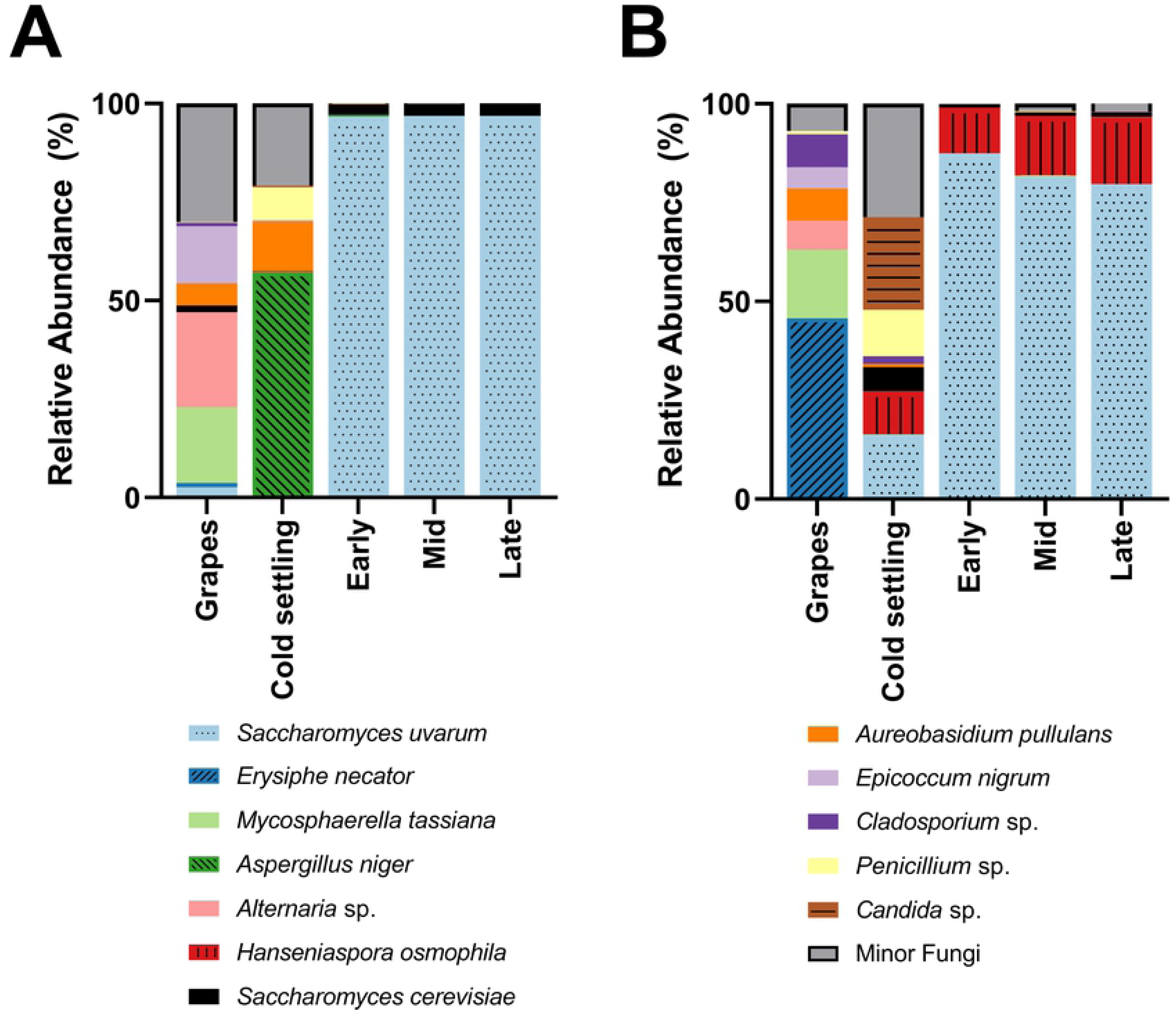
Fungal species abundance. Relative abundance of the dominant fungi present in grape samples taken in the vineyard (grapes), as well as at four stages of fermentation in the winery (cold settling, early, mid, and late), of Chardonnay sourced from (A) Vineyard 2 or (B) Vineyard 8. Vineyard 2 grape sample values are the means of five replicates, and Vineyard 8 grape sample values are the means of 6 replicates. All winery fermentation stages have three reported replicates, with the exception of the cold settling stage from the Vineyard 8 fermentations, which contained two. Relative abundance was calculated from 20,000 sequences per sample. Any fungal taxa that did not achieve at least 10% relative abundance in at least two samples were grouped into the Minor Fungi category, with one exception: *S. cerevisiae* did not achieve 10% in any one sample, but because of its importance during alcoholic fermentation it has been included here. For variation among samples please see S1 Table.

*A. pullulans* is a ubiquitous environmental yeast-like fungus that is commonly associated with grapes and vineyards [44,58,59], and was also the most common fungal species identified at the cold settling stage in must from Vineyard 8 two years prior to this current study [12]. *E. nigrum* is an endophytic fungus that has been previously identified in wine regions such as Italy, Portugal, and Spain [60–62]. Because both *A. pullulans* and *E. nigrum* are typically associated with vineyard environments, they have been proposed as potential biological control agents against grapevine pathogens [63]. *Alternaria* sp. are plant pathogens that are also commonly identified on grapes and in grape must [44,64,65]. *M. tassiana*, also known as *Davidiella tassiana*, is a common grape symbiont that has been previously isolated from vineyards [12,44,66,67], and has been identified as the most abundant fungal species isolated on grapes in one study [65]. The presence of *Apergillus* and *Penicillium* spp. in food, including grape must, has been observed previously [12,44,68]. These fungi have the potential to produce mycotoxins, which can be dangerous if consumed in large quantities [69,70]. However, low levels of mycotoxins are common in commercial wines [70], and Canadian wines typically have a lower concentration of mycotoxins than wines produced elsewhere in the world [71]. Furthermore, the process of alcoholic fermentation has been shown to reduce the presence of mycotoxins, through enzymatic conversion to a less toxic form and/or the adsorption to lees and subsequent removal from the wine [69,72]. *E. necator* (also called *Uncinula necator*) is a grapevine pathogen responsible for powdery mildew, and is found in all regions of the world where grapes are grown [73]. The dominant presence of this fungus in the grape samples of Vineyard 8 is of potential concern, because the presence of even small amounts can have a negative effect on the overall sensory profile of the wine [74]. However, the grapes sampled in this study were not visibly infected with powdery mildew, and the presence of *E. necator* seems to have been eliminated by the time the cold settling sample was taken (Fig 1B). In the Vineyard 8 cold settling sample we observed the presence of yeasts with fermentative potential such as *Candida* sp., *H. osmophila*, and *S. uvarum*. These fermentative yeasts were not identified in the Vineyard 2 cold settling sample, likely because the must from Vineyard 8 spent an extended amount of time in the settling tank before being transferred to barrels (where the cold settling samples were taken), and therefore was more susceptible to early winery-resident yeast exposure. Although *Candida* and *Hanseniaspora* spp. have been found in association with grapes in the vineyard [75], in this current study it seems more likely that the origin of these yeasts is the winery equipment that the grapes came into contact with, because neither *Candida* nor *Hanseniaspora* spp. were identified in the grape samples from Vineyard 8 (S1 Table). The grape samples from Vineyard 2 had a greater presence of *Saccharomyces* spp. than the grape samples from Vineyard 8: *S. cerevisiae* was present at ∼ 1.6% and *S. uvarum* at ∼ 2.6% in Vineyard 2, and at ∼ 0.05% and ∼ 0.03%, respectively, in Vineyard 8 (S1 Table). The abundance of *Saccharomyces* yeasts on healthy grapes in vineyards had been estimated at approximately 1/1000 yeast isolates [36], which is more in line with the results observed in Vineyard 8. The increased abundance of *Saccharomyces* yeasts in Vineyard 2 should be further investigated.

*S. uvarum* dominated the early, mid, and late stages of alcoholic fermentation in both vineyard treatments. In the Vineyard 2 fermentations, *S. uvarum* made up ∼ 96% of the relative abundance of these three stages, and *S. cerevisiae* made up ∼ 3%. In the Vineyard 8 fermentations, while *S. uvarum* still dominated with ∼ 80-87% of the relative abundance and *S. cerevisiae* was present at ∼ 1%, a third yeast, *H. osmophila*, was also present, maintaining ∼ 11-17% relative abundance through to the end of alcoholic fermentation. Originally, it was thought that *Hanseniaspora* spp. could not survive during the later stages of alcoholic fermentation due to low ethanol tolerance, because these yeasts could not be isolated from these stages using culture-dependent methods [76–78]. However, *H. osmophila* has been shown to have an ethanol tolerance of at least 9% [79], and the advent of culture-independent identification techniques such as high-throughput amplicon sequencing have allowed for the identification of all microorganisms present in wine fermentations, including those in a viable but not culturable (VBNC) state. Indeed, other studies that have employed similar culture-independent techniques to ours have identified *Hanseniaspora* spp. through to the end of alcoholic fermentation [44,57,80–82]. *H. osmophila* has been characterized as a glucophilic yeast [79,83,84], and may have contributed to the alcoholic fermentation in the fermentations from Vineyard 8.

*S. uvarum* was also found dominating Chardonnay fermentations at this same winery two years previously [12], suggesting that this is a winery-resident population, capable of overwintering and entering fermentations year after year. *S. uvarum* is usually found in association with white wine fermentations in cool-climate wine regions. It has been identified dominating such fermentations in wineries across Europe, including France [9,10], Hungary [11,30], Italy [85], and Slovakia [11]. *S. uvarum* is known to be cryotolerant [14,23], explaining its preference for low-temperature fermentations and cool-climate wine regions. The cellar in which the fermentations from this study were conducted is temperature-controlled and kept at 12 °C. This temperature is within a desirable growth range for *S. uvarum* [85,86], but is too low for *S. cerevisiae* to grow optimally [30,85]. To our knowledge, there is currently only one other winery (located in Alsace, France) that has been reported to have a local *S. uvarum* population dominating uninoculated fermentations across multiple vintages [9]. Interestingly, the Alsace cellar was also kept at 12 °C. It is possible that this temperature provides the optimal over-wintering conditions for *S. uvarum*, allowing it to out-compete *S. cerevisiae*, but more research is needed to investigate this.

### Saccharomyces uvarum strains

#### *S. uvarum* strain diversity

*S. uvarum* strain diversity was similarly high in the fermentations from both vineyards (Table 5), and this high diversity was maintained throughout alcoholic fermentation. The study conducted in 2015 at this same winery also reported high *S. uvarum* strain diversity that was established early and was maintained throughout alcoholic fermentation [12]. This result has also been observed with regards to *S. cerevisiae* strain diversity in uninoculated commercial fermentations [87,88].

**Table 5.**
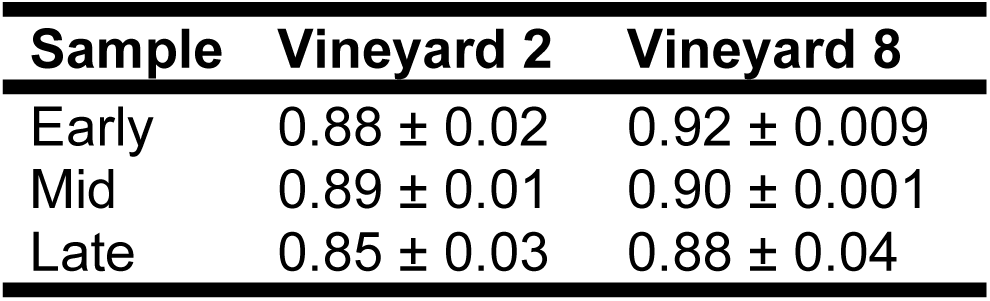
*Saccharomyces uvarum* strain diversity, measured as Simpson’s Index of Diversity (1 − *D*), of stainless steel barrel-fermented Chardonnay sourced from two different vineyards. Diversity ± SEM was measured from three stages of fermentation in the winery (early, mid, and late). All treatments and stages contained three replicates.

#### *S. uvarum* strain composition

A total of 106 unique *S. uvarum* strains were identified across the six fermentations sampled in this study (S2 Table). Two years previously, 150 unique *S. uvarum* strains were identified in fermentations at this same winery [12]; 66 of these strains were identified in both vintages. However, the previous study strain-typed 1,860 yeast isolates, in comparison to the 576 isolates strain-typed in this current study, so it is expected that fewer strains would be identified here. Other studies conducted to investigate *S. uvarum* strain abundance have identified far fewer strains (a maximum of 89 unique strains isolated from grapes, wine, and other environments), although it should be noted that fewer isolates were strain-typed in those studies (up to 114), potentially explaining this discrepancy [9–11,18,24,37,85,89].

Of the 106 strains identified in this study, only four were able to achieve ≥ 10% relative abundance in at least two samples (Fig 2). The other 102 strains, termed ‘minor strains,’ accounted for ∼ 50% of the relative abundance in the Vineyard 2 fermentations, and ∼ 65% of the relative abundance in the Vineyard 8 fermentations. The two most abundant strains in both vineyard treatments (‘2015 Strain 2’ and ‘2015 Strain 3’) were previously identified as dominant strains at the same winery two years previously, during the 2015 vintage [12]. The reoccurrence of ‘2015 Strain 2’ and ‘2015 Strain 3’ suggests that these strains have established themselves as persistent winery residents, capable of entering and dominating fermentations in multiple vintages. The other two dominant strains identified in these fermentations (‘2017 Strain 151’ and ‘2017 Strain 182’) were not isolated during the 2015 vintage, and were also not evenly distributed among the fermentations from the two vineyards (Fig 2). The ‘2017 Strain 151’ represented ∼ 7-17% of the relative abundance in the Vineyard 2 fermentations, and was only identified at ∼ 1% in a single sample in the Vineyard 8 fermentations. Meanwhile, the ‘2017 Strain 182’ represented ∼ 2-12% of the relative abundance in the Vineyard 8 fermentations, and was not identified at all in the Vineyard 2 fermentations. This difference in dominant strains could be attributed to a number of causes, including differences in acidity levels in the musts from the two different vineyards, or simply changes in the winery environment over time, since these fermentations did occur approximately two weeks apart. A significant correlation between grape must acidity and the dominance of specific *S. cerevisiae* strains has been previously observed [90], so it is understandable that a similar result might be expected with regards to *S. uvarum* strains.

**Fig 2.**
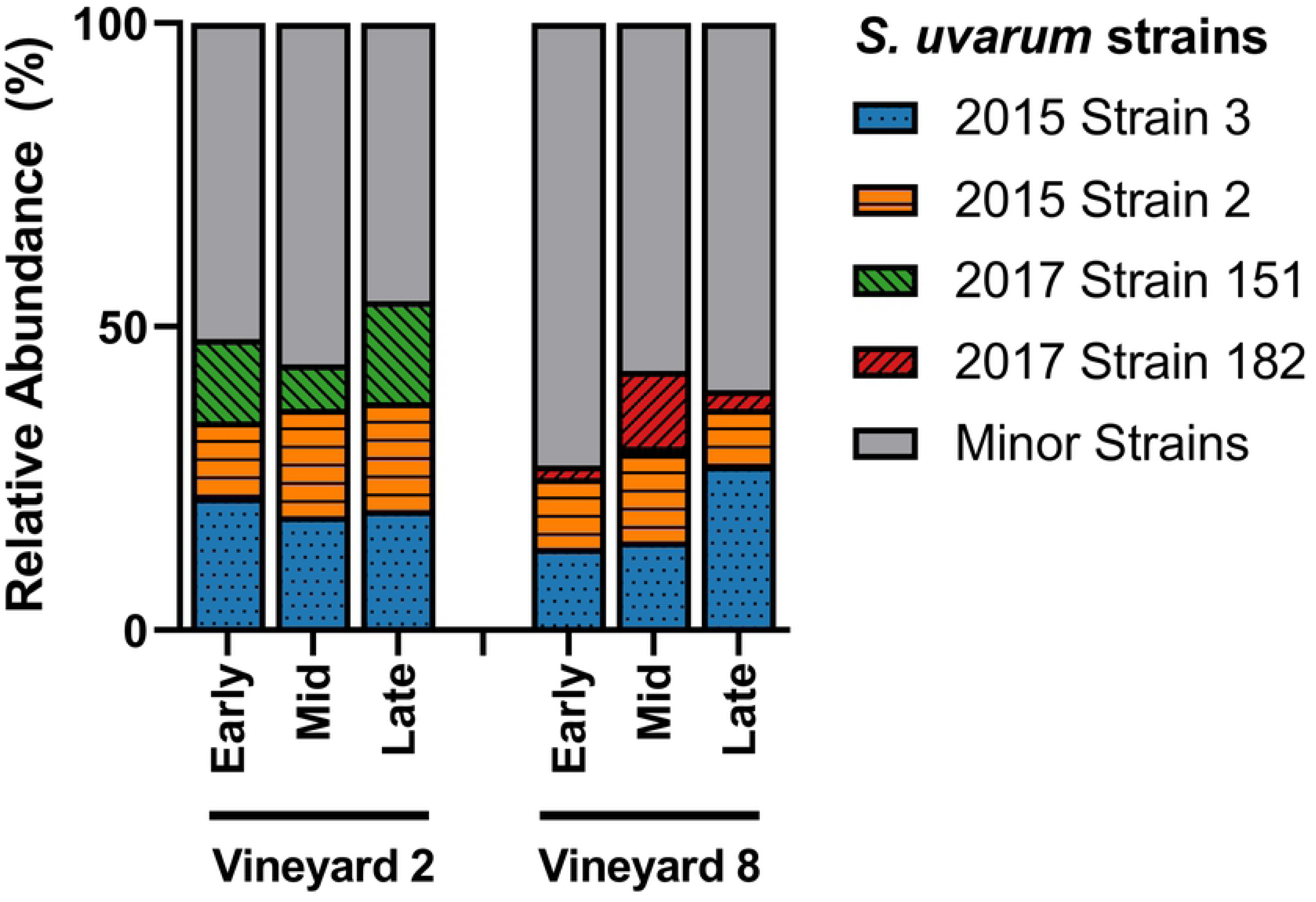
*S. uvarum* strain abundance. Relative abundance of the dominant *S. uvarum* strains present in three stages of fermentation (early, mid, and late) of Chardonnay sourced from two different vineyards (n = 3). Relative abundance was calculated from 32 *S. uvarum* yeast isolates per sample. Any strains that did not achieve ≥ 10% relative abundance in at least two samples were grouped into the Minor Strains category. For variation among samples please see S3 Table.

A principal coordinates analysis (PCoA) was also generated in order to visualize the differences in *S. uvarum* strain composition between the fermentations from the two vineyards while including all 106 strains (Fig 3). The PCoA ordination showed a clear separation between samples taken from the two vineyards. This result highlights the importance of analyzing not only the diversity of a sample but also the composition. In this study, as in previous studies of a similar nature [12,44], treatments have been found to have near-identical diversity values but completely distinct compositions. This is because composition considers not only the relative abundance of different strains, but also the identities of those strains, and can provide a more accurate summary of the differences observed among treatments. Previous research has identified that different strains of *S. uvarum* can produce wines with different sensory-active metabolite profiles, especially when fermented at lower temperatures [28,29], and it is therefore expected that different *S. uvarum* populations would contribute differently to the production of wine sensory attributes. However, this is still a new topic of investigation, and more research is needed in this area to determine whether the differences in secondary metabolite composition among wines fermented with different *S. uvarum* strains translate into detectable differences in the sensory profiles of the wines.

**Fig 3.**
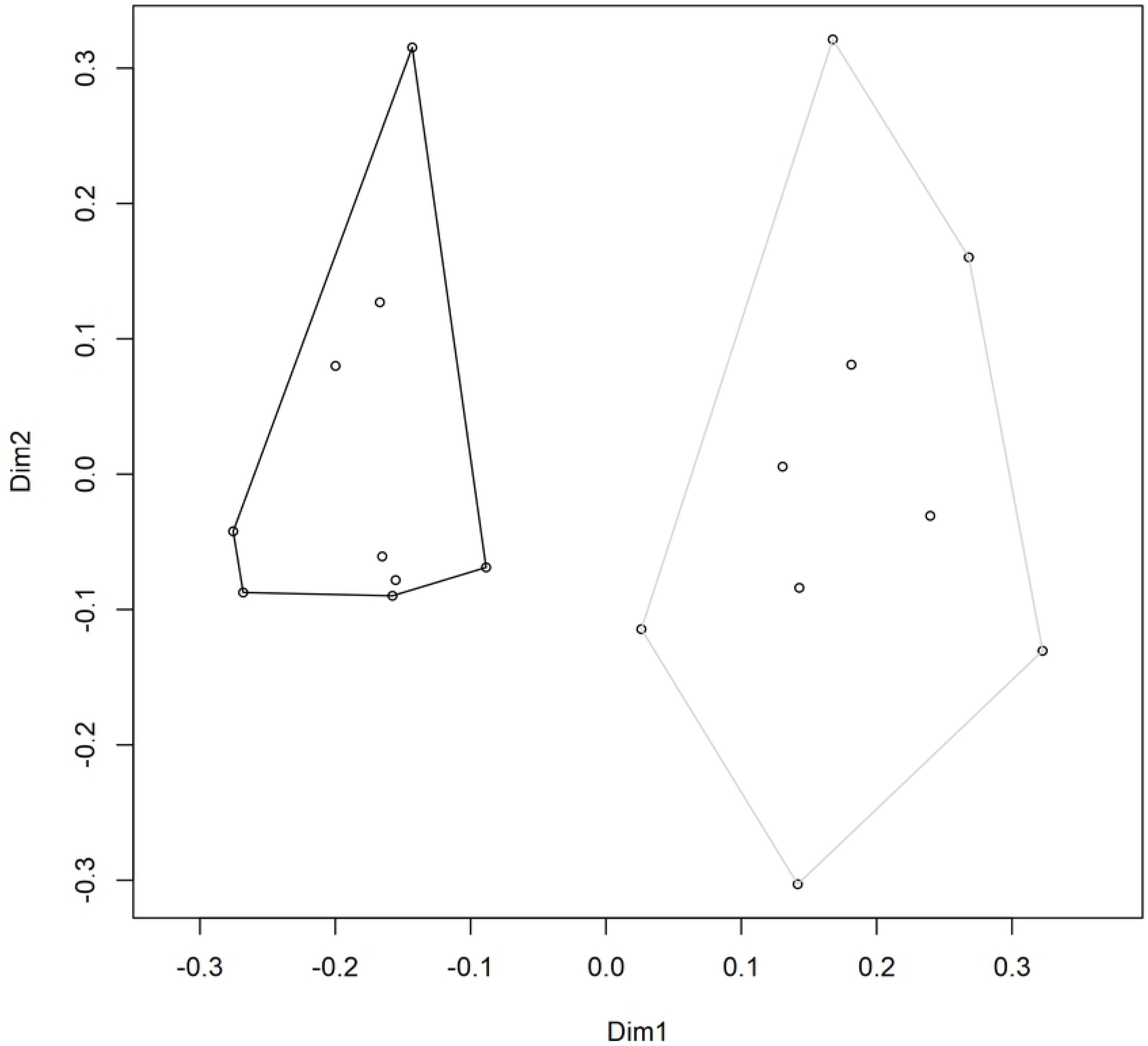
*S. uvarum* strain composition. Principal coordinates analysis (PCoA) ordination of the *S. uvarum* strain composition in Chardonnay wines sourced from two vineyards: Vineyard 2 (black) and Vineyard 8 (grey). Individual data points represent the composition of *S. uvarum* strains in a single sample (based on 32 yeast isolates per sample). Samples were taken at three stages of alcoholic fermentation, and each vineyard treatment contained three biological replicates, for a total of nine samples per vineyard. Dimension 1 (Dim1) explains 20.98% of variance, and Dimension 2 (Dim2) explains 13.07% of variance.

#### *S. uvarum* genetic diversity

Of the 106 strains identified in this study, 66 were also identified in the 2015 vintage at the same winery, including all four of the dominant strains from the 2015 vintage [12]. Additionally, we noted that 32 strains were unique to the Vineyard 2 fermentations, 40 strains were unique to the Vineyard 8 fermentations, and 34 strains were found in both the Vineyard 2 and Vineyard 8 fermentations. Although some minor strains were found to be unique to specific stages of fermentation, these strains were found in very low abundance: of the top 20 strains with the highest overall abundance, 18 were identified in all three stages of fermentation, suggesting that ethanol tolerance is not a major contributor to any differences in strain abundance observed.

Four minor strains were found to bear genetic similarity to international strains (within a Bruvo distance of 0.3). Seven isolates from the Vineyard 2 fermentations were genetically similar to the commercial strain Velluto BMV58^®^, despite this strain not being sold in Canada at the time of this study. Additionally, three isolates were genetically similar to the previously-sequenced Spanish strain CBS7001, one isolate was genetically similar to the French strain CBS8711, and one isolate was genetically similar to strain PYCC6860, which was isolated from oak trees on Hornby Island, in the same province of Canada as the winery from this study (although separated by hundreds of km). Local strains bearing genetic relatedness to international strains has been previously observed; some *S. uvarum* strains isolated in New Zealand were also found to bear significant nucleotide similarity with CBS7001 [24].

Eleven variable microsatellite loci were used to strain-type the *S. uvarum* isolates. The occurrence of heterozygosity in this study was higher than was observed at the same winery two years previously: 51.9% of the strains in this study contained at least one heterozygous locus, as compared to 42.7% in 2015 [12]. In both the 2017 and 2015 vintages at this winery, five was the average number of heterozygous loci. In this current study, 10 was the highest number of heterozygous loci identified in a single strain. This is the highest incidence of heterozygosity observed in an *S. uvarum* population to-date: previous studies have found heterozygous loci in 28.8% [37], 23.1% [24], and 0% [91] of *S. uvarum* strains isolated from grapes and wine. However, these studies did contain fewer isolates and analyzed fewer hypervariable microsatellite loci, which may explain some of the differences observed. The L9 microsatellite locus was the most variable by far, containing 18 different alleles in this study. The secondmost variable locus was L2, containing eight different alleles. The other loci contained either five alleles (L7 and L8), four alleles (L1, L3, L4, and NB9), or three alleles (NB1, NB4, and NB8).

A phylogenetic tree was created in order to visualize the genetic relatedness of the *S. uvarum* strains of significance from this study and the previous study conducted at this winery [12], as compared to 12 international *S. uvarum* strains isolated from winery and natural environments around the world (Table 2). Strains of significance are defined here as strains that represented > 1% of all the yeast isolates strain-typed in each vintage at this winery. In total, 43 strains were used to generate the unrooted phylogenetic tree (Fig 4). These strains were differentiated into four sub-populations using a *k*-means clustering algorithm, which aims to cluster the strains into the nearest groups so that the mean of each group has minimal variance. Group 1 contained nine strains, all of which were Okanagan strains originally isolated at the winery from this study in 2015. Group 2 contained 11 strains, including three Okanagan strains originally isolated in 2015, four Okanagan strains isolated in 2017, and four international strains, including one strain from the nearby Hornby Island. Group 3 contained 10 strains, including six Okanagan strains originally isolated in 2015 and four international strains, including one strain from the nearby Hornby Island. Group 4 contained 13 strains, including eight Okanagan strains originally isolated in 2015, one Okanagan strain isolated in 2017, and four international strains. The two strains isolated on Hornby Island were originally found associated with oak trees, not vineyard or winery environments, and appear to be natural *S. uvarum* isolates from Canada, because they lack the genetic introgressions from other *Saccharomyces* species that are seen in many European wine-associated *S. uvarum* strains [14]. One of these strains, PYCC6860, was also identified in one of the fermentations in this study. This finding is consistent with previous studies that report a mixing between natural and fermentation-derived *S. uvarum* strains in wine [37]. Groups 1, 2, and 3 all clustered separately within the tree, and while most of the strains in Group 4 clustered together, five strains from Group 4 were distributed throughout the tree (Fig 4). This means that the phylogenetic tree produced here shows clustering that does not follow the actual dendrogram assignments, suggesting that this is a panmictic population involving some random mating events, where the Okanagan strains identified in both 2015 and 2017 are not completely distinct subpopulations.

**Fig 4.**
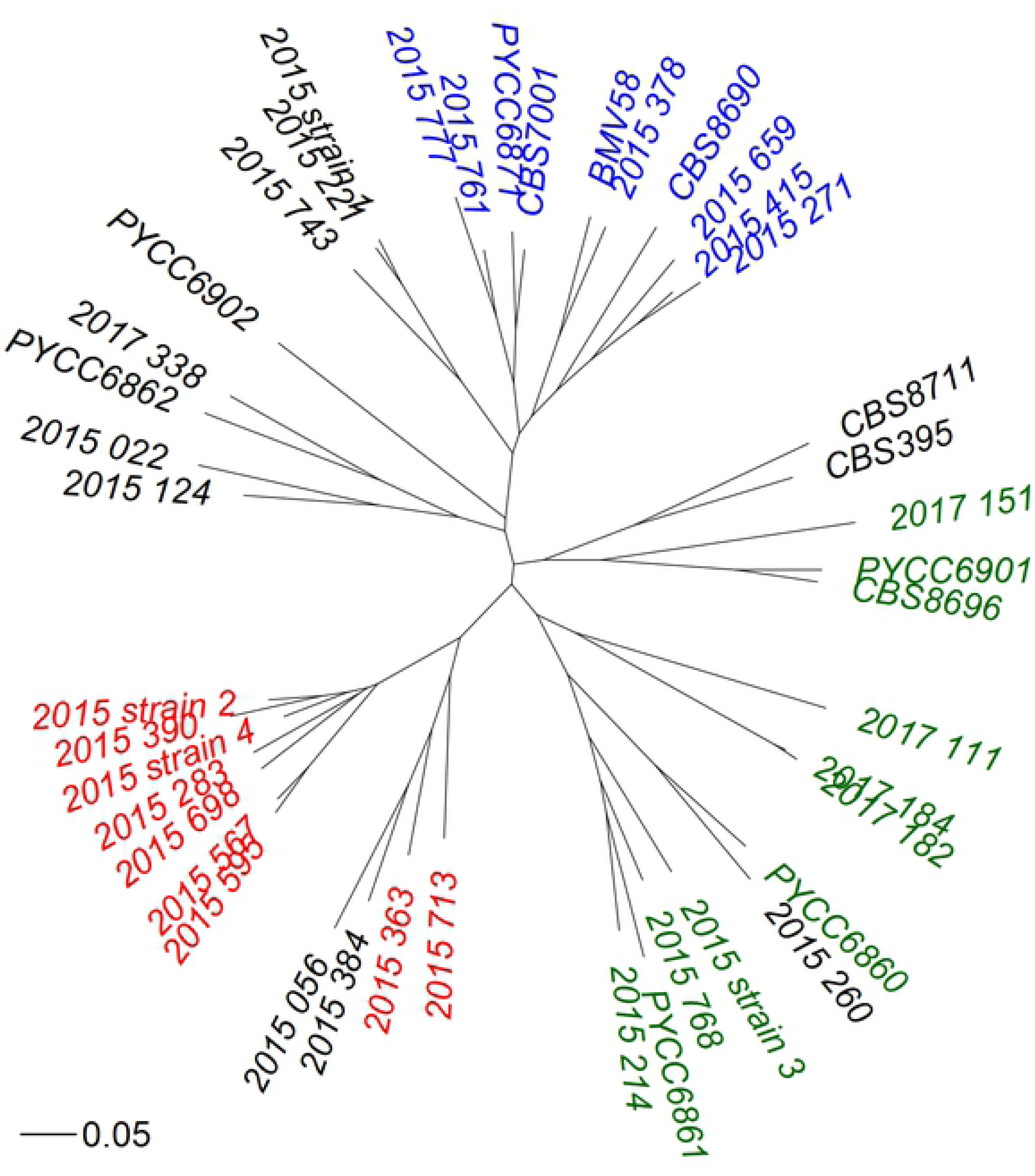
Phylogenetic tree of Okanagan and international *S. uvarum* strains. An unrooted phylogenetic tree using Bruvo distance, comparing the genetic relatedness of the *Saccharomyces uvarum* strains of significance identified Chardonnay fermentations in the Okanagan Valley (Canada) in both 2015 and 2017, as well as 12 selected *S. uvarum* strains isolated from around the world (see Table 2 for strain origins). A Bruvo distance of 0.05 is shown for scale. Strains of significance are defined here as strains that represented > 1% of all the yeast isolates strain-typed in each vintage. In total, 43 unique strains were compared, and were differentiated into four sub-populations using *k*-means clustering: Group 1 (Red), Group 2 (Green), Group 3 (Blue), and Group 4 (Black). The presence of some strains from Group 4 appearing throughout the tree suggests that this is a panmictic population involving random mating events.

The four dominant strains isolated in 2015 at the same winery (‘2015 Strain 1,’ ‘2015 Strain 2,’ ‘2015 Strain 3,’ and ‘2015 Strain 4’) were spread among Groups 1, 2, and 4. Meanwhile, the three of the four dominant strains isolated in 2017 (Fig 2) were found in Group 2, and the last was found in Group 1. Interestingly, none of the dominant strains from either 2015 or 2017 were found in Group 3. Only Group 1 contained no international strains, and this group also contained ‘2015 Strain 2,’ which was a dominant strain at this winery in both the 2015 and 2017 vintages. This suggests that the strains in Group 1 may belong to a uniquely Okanagan sub-population, but more research is needed to determine the natural origins of this population.

The only commercial *S. uvarum* strain included in the construction of this phylogenetic tree was Velluto^®^ BMV58 (Lallemand, Montreal, QC, Canada). Recently, one additional *S. uvarum* strain and one hybrid *S. uvarum* × *S. cerevisiae* strain have been commercially released: VitiFerm^TM^ Sauvage BIO and EnartisFerm^®^ ES U42, respectively. Some enological properties of *S. uvarum* have been studied, but more research is needed to allow winemakers to make informed decisions when selecting these non-traditional yeast strains for inoculation. *S. uvarum* produces lower levels of ethanol, acetic acid, and acetaldehyde, and higher levels of glycerol, succinic acid, malic acid, isoamyl alcohol, isobutanol, and ethyl acetate, as compared to *S. cerevisiae* [27–30]. Furthermore, due to its relatively lower production of ethanol, *S. uvarum* has been suggested as potential means of mitigating the effects of climate change on winemaking [92]. With a warming climate, many winemaking regions are beginning to produce very ripe grapes with high sugar contents, which, with traditional fermentation techniques, can result in wines with very high ethanol contents. If this trend continues, many wines produced in the future could contain alcohol concentrations above the legal regulations of some countries, since diluting grape must with water is not permitted in wine production. Using non-traditional yeasts such as *S. uvarum* or non-*Saccharomyces* species, which can metabolize grape sugars into compounds other than ethanol (such as glycerol), may help mitigate this issue by keeping ethanol production within permitted levels. The observed increase in the availability of commercial *S. uvarum* strains indicates that this is clearly an area of interest for winemakers, and our changing climate and consumer preferences requires investigation into new and creative methods of wine production, including the use of non-traditional yeasts such as *S. uvarum*, either alone or in combination with *S. cerevisiae*.

## Conclusions

This study investigated the fungal communities and *S. uvarum* populations present in uninoculated commercial Chardonnay fermentations of grapes that originated from two different vineyards. Differences in fungal community composition were observed, with *H. osmophila* representing a significant proportion of the fungal community in the fermentations from one vineyard, but not the other. However, in all of the fermentations, *S. uvarum* was the dominant yeast during the early, mid, and late stages of alcoholic fermentation. An investigation into the genetic diversity of the *S. uvarum* strains present in this study was conducted, and this population was found to be both highly diverse and genetically distinct from *S. uvarum* strains identified in other regions of the world. A total of 106 *S. uvarum* strains were identified in this study, but only four strains were able to play a dominant role in fermentation; two of these dominant strains were also identified as dominant strains at this same winery two years previously. The presence of persistent non-commercial strains, as well as the phylogenetic tree generated to compare Okanagan and international strains, provides evidence for an indigenous winery-resident *S. uvarum* population in the Okanagan Valley of Canada.

## Acknowledgements

The authors would like to thank the winemakers Darryl Brooker, Corrie Krehbiel, and Alexandra Haselich of Mission Hill Family Estate Winery for their assistance, guidance, and donation of fermentation samples. We also thank Britney Johnston and Mehrbod Estaki of the University of British Columbia for technical support and high-throughput amplicon sequencing data analysis support, respectively.

## Supporting Information Captions

**S1 Fig.**
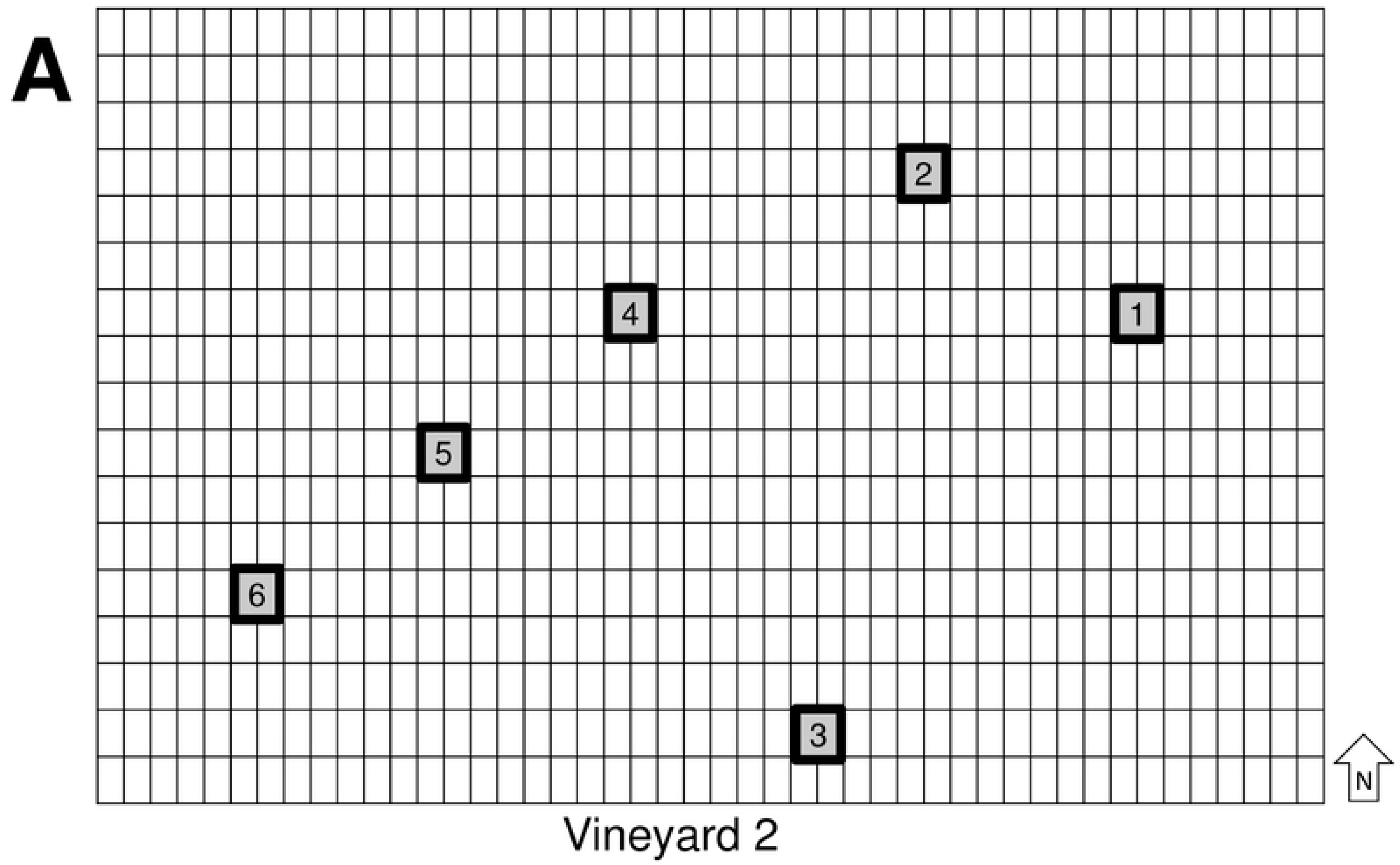
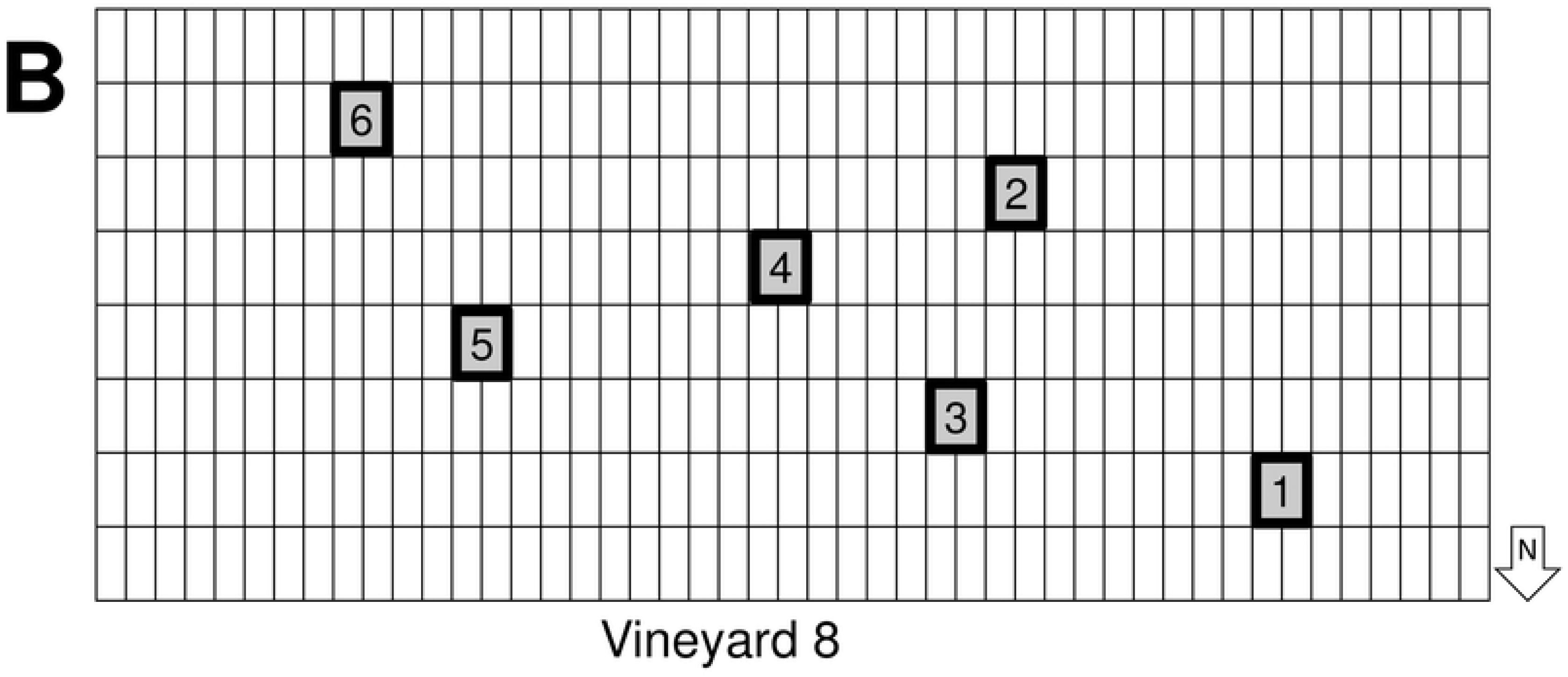
Vineyard sampling map. Sampling layouts for (A) Vineyard 2 and (B) Vineyard 8. Each of the two conjoined squares represent the generated site of collection for one sample, with the sample number also given at each sampling site. One sampling site contains approximately 15 vines, and two grape clusters were taken from each vine (one on either side of the row), for a total of 30 clusters per sampling site. The geographic orientation of each vineyard is indicated in the bottom right corner of each sampling map.

**S1 Table.** Fungal community composition of Chardonnay grapes, must, and fermenting wine sourced from two different vineyards, based on 20,000 sequences per sample and represented as percent (%) relative abundance. Samples were taken from grapes in the vineyard (G), and at four stages of fermentation in the winey: cold settling (C), early (E), mid (M), and late (L). Vineyard 2 grape sample values are the means ± SEM of five replicates, and Vineyard 8 grape sample values are the means ± SEM of 6 replicates. All winery fermentation stages have three reported replicates, with the exception of the cold settling stage from the Vineyard 8 fermentations, which contained two. Sequences were identified to the species level unless otherwise indicated. Fungal species that represented less than 10% of the relative abundance in at least two samples were grouped into the Minor Fungi category. One exception was *Saccharomyces cerevisiae*, which never reached 10% relative abundance in any sample but is included in this table because of its importance during alcoholic fermentation. The last two columns indicate positive (Pos) and negative (Neg) controls. For the raw data containing all the fungi identified in this study (including minor fungi), please visit https://osf.io/j7rx8/.

**S2 Table**. Microsatellite identities of the representative multilocus genotypes (MLGs) of *Saccharomyces uvarum* strains isolated from stainless steel barrel-fermented Chardonnay at Canadian winery during the 2017 vintage. Allele sizes for allele 1 (A1) and allele 2 (A2) are shown for each of the 11 microsatellite loci analyzed. Strains with the prefix “2017” were isolated exclusively during the 2017 vintage. Strains with the prefix “2015” were previously isolated and characterized during the 2015 vintage at the same winery, and were also isolated during the 2017 vintage. Strains without a vintage prefix are those that belong to global yeast databases (see S4 Table).

**S3 Table.** *Saccharomyces uvarum* population composition of Chardonnay grapes, must, and fermenting wine sourced from two different vineyards, based on 32 yeast isolates per sample. Samples were taken at three stages of fermentation: early (E), mid (M), and late (L). Values are the means ± SEM (n = 3). Strains that represented less than 10% of the relative abundance in at least two samples were grouped into the Minor Strains category. For the raw data, containing all the *S. uvarum* strains identified in this study (including minor strains), please visit https://osf.io/j7rx8/.

**S4 Table**. Microsatellite identities of the multilocus genotypes (MLGs) of *Saccharomyces uvarum* strains obtained from global yeast databases. Allele sizes for allele 1 (A1) and allele 2 (A2) are shown for each of the 11 microsatellite loci.

## References

1. Suárez-Lepe JA, Morata A. New trends in yeast selection for winemaking. Trends Food Sci Technol. 2012;23: 39–50. doi:10.1016/j.tifs.2011.08.005

2. Albertin W, Zimmer A, Miot-Sertier C, Bernard M, Coulon J, Moine V, et al. Combined effect of the *Saccharomyces cerevisiae* lag phase and the non-*Saccharomyces* consortium to enhance wine fruitiness and complexity. Appl Microbiol Biotechnol. 2017;101: 7603–7620. doi:10.1007/s00253-017-8492-1

3. Capozzi V, Garofalo C, Chiriatti MA, Grieco F, Spano G. Microbial terroir and food innovation: The case of yeast biodiversity in wine. Microbiol Res. 2015;181: 75– 83. doi:10.1016/j.micres.2015.10.005

4. Knight S, Klaere S, Fedrizzi B, Goddard MR. Regional microbial signatures positively correlate with differential wine phenotypes: Evidence for a microbial aspect to terroir. Sci Rep. 2015;5: 14233. doi:10.1038/srep14233

5. Scholl CM, Morgan SC, Stone ML, Tantikachornkiat M, Neuner M, Durall DM. Composition of *Saccharomyces cerevisiae* strains in spontaneous fermentations of Pinot Noir and Chardonnay. Aust J Grape Wine Res. 2016;22: 384–390. doi:10.1111/ajgw.12221

6. Ciani M, Mannazzu I, Marinangeli P, Clementi F, Martini A. Contribution of winery-resident *Saccharomyces cerevisiae* strains to spontaneous grape must fermentation. Antonie Van Leeuwenhoek. 2004;85: 159–164.

7. Santamaría P, López R, López E, Garijo P, Gutiérrez AR. Permanence of yeast inocula in the winery ecosystem and presence in spontaneous fermentations. Eur Food Res Technol. 2008;227: 1563–1567. doi:10.1007/s00217-008-0855-5

8. Blanco P, Orriols I, Losada a. Survival of commercial yeasts in the winery environment and their prevalence during spontaneous fermentations. J Ind Microbiol Biotechnol. 2011;38: 235–239. doi:10.1007/s10295-010-0818-2

9. Demuyter C, Lollier M, Legras J-L, Le Jeune C. Predominance of *Saccharomyces uvarum* during spontaneous alcoholic fermentation, for three consecutive years, in an Alsatian winery. J Appl Microbiol. 2004;97: 1140–1148. doi:10.1111/j.1365-2672.2004.02394.x

10. Naumov GI, Masneuf I, Naumova ES, Aigle M, Dubourdieu D. Association of *Saccharomyces bayanus* var. *uvarum* with some French wines: Genetic analysis of yeast populations. Res Microbiol. 2000;151: 683–691. doi:10.1016/S0923-2508(00)90131-1

11. Naumov GI, Naumova ES, Antunovics Z, Sipiczki M. *Saccharomyces bayanus* var. *uvarum* in Tokaj wine-making of Slovakia and Hungary. Appl Microbiol Biotechnol. 2002;59: 727–730. doi:10.1007/s00253-002-1077-6

12. Morgan SC, McCarthy GC, Watters BS, Tantikachornkiat M, Zigg I, Cliff MA, et al. Effect of sulfite addition and *pied de cuve* inoculation on the microbial communities and sensory profiles of Chardonnay wines: dominance of indigenous *Saccharomyces uvarum* at a commercial winery. FEMS Yeast Res. 2019;19: foz049. doi:10.1093/femsyr/foz049

13. Borneman AR, Pretorius IS. Genomic insights into the *Saccharomyces sensu stricto* complex. Genetics. 2015;199: 281–291. doi:10.1534/genetics.114.173633

14. Almeida P, Gonçalves C, Teixeira S, Libkind D, Bontrager M, Masneuf-Pomarède I, et al. A Gondwanan imprint on global diversity and domestication of wine and cider yeast *Saccharomyces uvarum*. Nat Commun. 2014;5: 4044. doi:10.1038/ncomms5044

15. Libkind D, Hittinger C, Valério E, Gonçalves C, Dover J, Johnston M, et al. Microbe domestication and the identification of the wild genetic stock of lager-brewing yeast. Proc Natl Acad Sci. 2011;108: 14539–14544. doi:10.1073/pnas.1105430108

16. Sampaio JP, Gonçalves P. Natural populations of *Saccharomyces kudriavzevii* in Portugal are associated with oak bark and are sympatric with *S. cerevisiae* and *S. paradoxus*. Appl Environ Microbiol. 2008;74: 2144–2152. doi:10.1128/AEM.02396-07

17. Naumov GI, Nguyen H-V, Naumova ES, Michel A, Aigle M, Gaillardin C. Genetic identification of *Saccharomyces bayanus* var. *uvarum*, a cider-fermenting yeast. Int J Food Microbiol. 2001.

18. Masneuf-Pomarede I, Le Jeune C, Durrens P, Lollier M, Aigle M, Dubourdieu D. Molecular typing of wine yeast strains *Saccharomyces bayanus* var. *uvarum* using microsatellite markers. Syst Appl Microbiol. 2007;30: 75–82. doi:10.1016/j.syapm.2006.02.006

19. Coton E, Coton M, Levert D, Casaregola S, Sohier D. Yeast ecology in French cider and black olive natural fermentations. Int J Food Microbiol. 2006;108: 130– 135.

20. Suárez Valles B, Pando Bedriñana R, Fernández Tascón N, Querol Simón A, Rodríguez Madrera R. Yeast species associated with the spontaneous fermentation of cider. Food Microbiol. 2007;24: 25–31. doi:10.1016/j.fm.2006.04.001

21. Fernández-Espinar MT, López V, Ramón D, Bartra E, Querol A. Study of the authenticity of commercial wine yeast strains by molecular techniques. Int J Food Microbiol. 2001;70: 1–10. doi:10.1016/S0168-1605(01)00502-5

22. Nguyen HV, Gaillardin C. Evolutionary relationships between the former species *Saccharomyces uvarum* and the hybrids *Saccharomyces bayanus* and *Saccharomyces pastorianus*; reinstatement of *Saccharomyces uvarum* (Beijerinck) as a distinct species. FEMS Yeast Res. 2005;5: 471–483. doi:10.1016/j.femsyr.2004.12.004

23. Rainieri S, Zambonelli C, Hallsworth JE, Pulvirenti A, Giudici P. *Saccharomyces uvarum*, a distinct group within *Saccharomyces sensu stricto*. FEMS Microbiol Lett. 1999;177: 177–185. doi:10.1111/j.1574-6968.1999.tb13729.x

24. Zhang H, Richards KD, Wilson S, Lee SA, Sheehan H, Roncoroni M, et al. Genetic characterization of strains of *Saccharomyces uvarum* from New Zealand wineries. Food Microbiol. 2015;46: 92–99. doi:10.1016/j.fm.2014.07.016

25. Rodríguez ME, Pérez-Través L, Sangorrín MP, Barrio E, Lopes CA. *Saccharomyces eubayanus* and *Saccharomyces uvarum* associated with the fermentation of *Araucaria araucana* seeds in Patagonia. FEMS Yeast Res. 2014;14: 948–965. doi:10.1111/1567-1364.12183

26. Rodríguez M, Pérez-Través L, Sangorrín M, Barrio E, Querol A, Lopes C. *Saccharomyces uvarum* is responsible for the traditional fermentation of apple *Chicha* in Patagonia. FEMS Yeast Res. 2017;17: fow109. doi:10.1093/femsyr/fow109

27. Castellari L, Ferruzzi M, Magrini A, Giudici P, Passarelli P, Zambonelli C. Unbalanced wine fermentation by cryotolerant vs. non-cryotolerant *Saccharomyces* strains. Vitis. 1994;33: 49–52.

28. Gamero A, Tronchoni J, Querol A, Belloch C. Production of aroma compounds by cryotolerant *Saccharomyces* species and hybrids at low and moderate fermentation temperatures. J Appl Microbiol. 2013;114: 1405–1414. doi:10.1111/jam.12126

29. Magyar I, Tóth T. Comparative evaluation of some oenological properties in wine strains of *Candida stellata*, *Candida zemplinina*, *Saccharomyces uvarum* and *Saccharomyces cerevisiae*. Food Microbiol. 2011;28: 94–100. doi:10.1016/j.fm.2010.08.011

30. Sipiczki M, Romano P, Lipani G, Miklos I, Antunovics Z. Analysis of yeasts derived from natural fermentation in a Tokaj winery. Antonie van Leeuwenhoek, Int J Gen Mol Microbiol. 2001;79: 97–105. doi:10.1023/A:1010249408975

31. Deed RC, Fedrizzi B, Gardner RC. Influence of fermentation temperature, yeast strain, and grape juice on the aroma chemistry and sensory profile of Sauvignon blanc wines. J Agric Food Chem. 2017;65: 8902–8912. doi:10.1021/acs.jafc.7b03229

32. Castelluci F. Microbiological analysis of wines and musts Method OIV-MA-AS4-01 Type IV Method. Compendium of International Methods of Analysis - OIV. OIV; 2010. pp. 1–32.

33. Houck L. Use of biphenyl for reducing Penicillium decay of stored citrus. Agric Res Serv. 1971.

34. Martiniuk JT, Pacheco B, Russell G, Tong S, Backstrom I, Measday V. Impact of commercial strain use on *Saccharomyces cerevisiae* population structure and dynamics in Pinot noir vineyards and spontaneous fermentations of a Canadian winery. PLoS One. 2016;11: e0160259. doi:10.1371/journal.pone.0160259

35. Zott K, Miot-Sertier C, Claisse O, Lonvaud-Funel A, Masneuf-Pomarede I. Dynamics and diversity of non-*Saccharomyces* yeasts during the early stages in winemaking. Int J Food Microbiol. 2008;125: 197–203. doi:10.1016/j.ijfoodmicro.2008.04.001

36. Mortimer R, Polsinelli M. On the origins of wine yeast. Res Microbiol. 1999;150: 199–204. doi:10.1016/S0923-2508(99)80036-9

37. Masneuf-Pomarede I, Salin F, Börlin M, Coton E, Coton M, Le Jeune C, et al. Microsatellite analysis of *Saccharomyces uvarum* diversity. FEMS Yeast Res. 2016;16: fow002. doi:10.1093/femsyr/fow002

38. Bruvo R, Michiels NK, D’Souza TG, Schulenburg H. A simple method for the calculation of microsatellite genotype distances irrespective of ploidy level. Mol Ecol. 2004;13: 2101–2106. doi:10.1111/j.1365-294X.2004.02209.x

39. Kamvar ZN, Brooks JC, Grünwald NJ. Novel R tools for analysis of genome-wide population genetic data with emphasis on clonality. Front Genet. 2015;6: 208. doi:10.3389/fgene.2015.00208

40. Kamvar ZN, Tabima JF, Grünwald NJ. Poppr: An R package for genetic analysis of populations with clonal, partially clonal, and/or sexual reproduction. PeerJ. 2014;2: e281. doi:10.7717/peerj.281

41. Paradis E, Schliep K. ape 5.0: an environment for modern phylogenetics and evolutionary analyses in R. Schwartz R, editor. Bioinformatics. 2019;35: 526–528. doi:10.1093/bioinformatics/bty633

42. Jombart T, Devillard S, Balloux F. Discriminant analysis of principal components: a new method for the analysis of genetically structured populations. BMC Genet. 2010;11: 94. doi:10.1186/1471-2156-11-94

43. Zott K, Claisse O, Lucas P, Coulon J, Lonvaud-Funel A, Masneuf-Pomarede I. Characterization of the yeast ecosystem in grape must and wine using real-time PCR. Food Microbiol. 2010;27: 559–567. doi:10.1016/J.FM.2010.01.006

44. Morgan SC, Tantikachornkiat M, Scholl CM, Benson NL, Cliff MA, Durall DM. The effect of sulfur dioxide addition at crush on the fungal and bacterial communities and the sensory attributes of Pinot gris wines. Int J Food Microbiol. 2019;290: 1– 14. doi:10.1016/j.ijfoodmicro.2018.09.020

45. Bokulich NA, Mills DA. Improved selection of Internal Transcribed Spacer-specific primers enables quantitative, ultra-high-throughput profiling of fungal communities. Appl Environ Microbiol. 2013;79: 2519–2526. doi:10.1128/AEM.03870-12

46. Bolyen E, Rideout JR, Dillon MR, Bokulich NA, Abnet CC, Al-Ghalith GA, et al. Reproducible, interactive, scalable and extensible microbiome data science using QIIME 2. Nat Biotechnol. 2019;7. doi:10.1038/s41587-019-0209-9

47. Callahan BJ, McMurdie PJ, Rosen MJ, Han AW, Johnson AJA, Holmes SP. DADA2: High resolution sample inference from Illumina amplicon data. Nat Methods. 2016;13: 581–583. doi:10.1038/nmeth.3869.DADA2

48. Morgan M, Anders S, Lawrence M, Aboyoun P, Pagès H, Gentleman R. ShortRead: A bioconductor package for input, quality assessment and exploration of high-throughput sequence data. Bioinformatics. 2009;25: 2607–2608. doi:10.1093/bioinformatics/btp450

49. Martin M. Cutadapt removes adapter sequences from high-throughput sequencing reads. EMBnet.journal. 2011;17: 10–12. doi:10.14806/ej.17.1.200

50. Blaalid R, Kumar S, Nilsson RH, Abarenkov K, Kirk PM, Kauserud H. ITS1 versus ITS2 as DNA metabarcodes for fungi. Mol Ecol Resour. 2013;13: 218–224. doi:10.1111/1755-0998.12065

51. Feibelman T, Bayman P, Cibula WG. Length variation in the internal transcribed spacer of ribosomal DNA in chanterelles. Mycol Res. 1994;98: 614–618. doi:10.1016/S0953-7562(09)80407-3

52. Katoh K, Standley DM. MAFFT multiple sequence alignment software version 7: Improvements in performance and usability. Mol Biol Evol. 2013;30: 772–780. doi:10.1093/molbev/mst010

53. Price MN, Dehal PS, Arkin AP. FastTree 2 – Approximately Maximum-Likelihood Trees for Large Alignments. Poon AFY, editor. PLoS One. 2010;5: e9490. doi:10.1371/journal.pone.0009490

54. Kõljalg U, Nilsson RH, Abarenkov K, Tedersoo L, Taylor AFS, Bahram M, et al. Towards a unified paradigm for sequence-based identification of fungi. Mol Ecol. 2013;22: 5271–5277. doi:10.1111/mec.12481

55. Schütz M, Gafner J. Lower fructose uptake capacity of genetically characterized strains of Saccharomyces bayanus compared to strains of Saccharomyces cerevisiae: a likely cause of reduced alcoholic fermentation activity. Am J Enol Vitic. 1995;46: 175–180.

56. Amerine MA, Roessler EB. Wines: Their Sensory Evaluation. New York: W.H.Freeman & Co Ltd; 1983.

57. Wang C, Mas A, Esteve-Zarzoso B. The interaction between *Saccharomyces cerevisiae* and non-*Saccharomyces* yeast during alcoholic fermentation is species and strain specific. Front Microbiol. 2016;7: 1–11. doi:10.3389/fmicb.2016.00502

58. Varela C, Borneman AR. Yeasts found in vineyards and wineries. Yeast. 2017;34: 111–128. doi:10.1002/yea.3219

59. Takahashi M, Ohta T, Masaki K, Mizuno A, Goto-Yamamoto N. Evaluation of microbial diversity in sulfite-added and sulfite-free wine by culture-dependent and -independent methods. J Biosci Bioeng. 2014;117: 569–75. doi:10.1016/j.jbiosc.2013.10.012

60. González V, Tello ML. The endophytic mycota associated with *Vitis vinifera* in central Spain. Fungal Divers. 2011;47: 29–42. doi:10.1007/s13225-010-0073-x

61. Pancher M, Ceol M, Corneo PE, Longa CMO, Yousaf S, Pertot I, et al. Fungal endophytic communities in grapevines (*Vitis vinifera* L.) Respond to crop management. Appl Environ Microbiol. 2012;78: 4308–4317. doi:10.1128/AEM.07655-11

62. Varanda CMR, Oliveira M, Materatski P, Landum M, Clara MIE, Félix M do R. Fungal endophytic communities associated to the phyllosphere of grapevine cultivars under different types of management. Fungal Biol. 2016;120: 1525– 1536. doi:10.1016/j.funbio.2016.08.002

63. Martini M, Musetti R, Grisan S, Polizzotto R, Borselli S, Pavan F, et al. DNA-dependent detection of the grapevine fungal endophytes *Aureobasidium pullulans* and *Epicoccum nigrum*. Plant Dis. 2009;93: 993–998. doi:10.1094/PDIS-93-10-0993

64. Kecskeméti E, Berkelmann-Löhnertz B, Reineke A. Are epiphytic microbial communities in the carposphere of ripening grape clusters (*Vitis vinifera* L.) different between conventional, organic, and biodynamic grapes? PLoS One. 2016;11. doi:10.1371/journal.pone.0160852

65. Castañeda LE, Miura T, Sánchez R, Barbosa O. Effects of agricultural management on phyllosphere fungal diversity in vineyards and the association with adjacent native forests. PeerJ. 2018;6: e5715. doi:10.7717/peerj.5715

66. Bokulich NA, Thorngate JH, Richardson PM, Mills DA. Microbial biogeography of wine grapes is conditioned by cultivar, vintage, and climate. Proc Natl Acad Sci. 2014;111: E139–E148. doi:10.1073/pnas.1317377110

67. Setati ME, Jacobson D, Bauer FF. Sequence-based analysis of the *Vitis vinifera* L. cv Cabernet sauvignon grape must mycobiome in three South African vineyards employing distinct agronomic systems. Front Microbiol. 2015;6: 1358. doi:10.3389/fmicb.2015.01358

68. Freire L, Passamani FRF, Thomas AB, Nassur R de CMR, Silva LM, Paschoal FN, et al. Influence of physical and chemical characteristics of wine grapes on the incidence of *Penicillium* and *Aspergillus* fungi in grapes and ochratoxin A in wines. Int J Food Microbiol. 2017;241: 181–190. doi:10.1016/j.ijfoodmicro.2016.10.027

69. Inoue T, Nagatomi Y, Uyama A, Mochizuki N. Degradation of aflatoxin B1 during the fermentation of alcoholic beverages. Toxins (Basel). 2013;5: 1219–1229. doi:10.3390/toxins5071219

70. Mateo R, Medina Á, Mateo EM, Mateo F, Jiménez M. An overview of ochratoxin A in beer and wine. Int J Food Microbiol. 2007;119: 79–83. doi:10.1016/j.ijfoodmicro.2007.07.029

71. Soleas GJ, Yan J, Goldberg DM. Assay of ochratoxin A in wine and beer by high-pressure liquid chromatography photodiode array and gas chromatography mass selective detection. J Agric Food Chem. 2001;49: 2733–2740. doi:10.1021/jf0100651

72. Petruzzi L, Sinigaglia M, Corbo MR, Campaniello D, Speranza B, Bevilacqua A. Decontamination of ochratoxin A by yeasts: Possible approaches and factors leading to toxin removal in wine. Appl Microbiol Biotechnol. 2014;98: 6555–6567. doi:10.1007/s00253-014-5814-4

73. Brewer MT, Milgroom MG. Phylogeography and population structure of the grape powdery mildew fungus, *Erysiphe necator*, from diverse *Vitis* species. BMC Evol Biol. 2010;10: 268. doi:10.1186/1471-2148-10-268

74. Gadoury DM, Seem RC, Wilcox WF, Henick-Kling T, Conterno L, Day A, et al. Effects of diffuse colonization of grape berries by *Uncinula necator* on bunch rots, berry microflora, and juice and wine quality. Phytopathology. 2007;97: 1356– 1365. doi:10.1094/PHYTO-97-10-1356

75. Barata A, Malfeito-Ferreira M, Loureiro V. The microbial ecology of wine grape berries. Int J Food Microbiol. 2012;153: 243–259. doi:10.1016/j.ijfoodmicro.2011.11.025

76. Combina M, Elía A, Mercado L, Catania C, Ganga A, Martinez C. Dynamics of indigenous yeast populations during spontaneous fermentation of wines from Mendoza, Argentina. Int J Food Microbiol. 2005;99: 237–243. doi:10.1016/j.ijfoodmicro.2004.08.017

77. Constantí M, Poblet M, Arola L, Mas A, Guillamón JM. Analysis of yeast population during alcoholic fermentation in a newly established winery. Am J Enol Vitic. 1997;48: 339–343.

78. Ocón E, Gutiérrez a. R, Garijo P, Tenorio C, López I, López R, et al. Quantitative and qualitative analysis of non-*Saccharomyces* yeasts in spontaneous alcoholic fermentations. Eur Food Res Technol. 2010;230: 885–891. doi:10.1007/s00217-010-1233-7

79. Granchi L, Ganucci D, Messini A, Vincenzini M. Oenological properties of *Hanseniaspora osmophila* and *Kloeckera corticis* from wines produced by spontaneous fermentations of normal and dried grapes. FEMS Yeast Res. 2002;2: 403–407. doi:10.1111/j.1567-1364.2002.tb00110.x

80. Pateraki C, Paramithiotis S, Doulgeraki AI, Kallithraka S, Kotseridis Y, Drosinos EH. Effect of sulfur dioxide addition in wild yeast population dynamics and polyphenolic composition during spontaneous red wine fermentation from *Vitis vinifera* cultivar Agiorgitiko. Eur Food Res Technol. 2014;239: 1067–1075. doi:10.1007/s00217-014-2303-z

81. Stefanini I, Albanese D, Cavazza A, Franciosi E, De Filippo C, Donati C, et al. Dynamic changes in microbiota and mycobiota during spontaneous “Vino Santo Trentino” fermentation. Microb Biotechnol. 2016;9: 195–208. doi:10.1111/1751-7915.12337

82. Bokulich NA, Swadener M, Sakamoto K, Mills DA, Bisson LF. Sulfur dioxide treatment alters wine microbial diversity and fermentation progression in a dose-dependent fashion. Am J Enol Vitic. 2014;66: 1–21. doi:10.5344/ajev.2014.14096

83. Viana F, Gil J V., Genovés S, Vallés S, Manzanares P. Rational selection of non-*Saccharomyces* wine yeasts for mixed starters based on ester formation and enological traits. Food Microbiol. 2008;25: 778–785. doi:10.1016/j.fm.2008.04.015

84. Ciani M, Comitini F, Mannazzu I, Domizio P. Controlled mixed culture fermentation: A new perspective on the use of non-*Saccharomyces* yeasts in winemaking. FEMS Yeast Res. 2010;10: 123–133. doi:10.1111/j.1567-1364.2009.00579.x

85. Dellaglio F, Zapparoli G, Malacrinò P, Suzzi G, Torriani S. *Saccharomyces bayanus* var. *uvarum* and *Saccharomyces cerevisiae* succession during spontaneous fermentations of Recioto and Amarone wines. Ann Microbiol. 2003.

86. Csoma H, Zakany N, Capece A, Romano P, Sipiczki M. Biological diversity of *Saccharomyces* yeasts of spontaneously fermenting wines in four wine regions: Comparative genotypic and phenotypic analysis. Int J Food Microbiol. 2010;140: 239–248. doi:10.1016/j.ijfoodmicro.2010.03.024

87. Morgan SC, Scholl CM, Benson NL, Stone ML, Durall DM. Sulfur dioxide addition at crush alters *Saccharomyces cerevisiae* strain composition in spontaneous fermentations at two Canadian wineries. Int J Food Microbiol. 2017;244: 96–102. doi:10.1016/j.ijfoodmicro.2016.12.025

88. Sabate J, Cano J, Querol A, Guillamon JM. Diversity of *Saccharomyces* strains in wine fermentations: analysis for two consecutive years. Lett Appl Microbiol. 1998;26: 452–455. doi:10.1046/j.1472-765X.1998.00369.x

89. Tosi E, Azzolini M, Guzzo F, Zapparoli G. Evidence of different fermentation behaviours of two indigenous strains of *Saccharomyces cerevisiae* and *Saccharomyces uvarum* isolated from Amarone wine. J Appl Microbiol. 2009;107: 210–218. doi:10.1111/j.1365-2672.2009.04196.x

90. Blanco P, Mirás-Avalos JM, Orriols I. Effect of must characteristics on the diversity of *Saccharomyces* strains and their prevalence in spontaneous fermentations. J Appl Microbiol. 2012;112: 936–944. doi:10.1111/j.1365-2672.2012.05278.x

91. Masneuf-Pomarède I, Le Jeune C, Durrens P, Lollier M, Aigle M, Dubourdieu D. Molecular typing of wine yeast strains Saccharomyces bayanus var. uvarum using microsatellite markers. Syst Appl Microbiol. 2007;30: 75–82. doi:10.1016/J.SYAPM.2006.02.006

92. Alonso-del-Real J, Lairón-Peris M, Barrio E, Querol A. Effect of temperature on the prevalence of *Saccharomyces* non *cerevisiae* species against a *S. cerevisiae* wine strain in wine fermentation: competition, physiological fitness, and influence in final wine composition. Front Microbiol. 2017;8: 1–15. doi:10.3389/fmicb.2017.00150

